# A brainstem-thalamic circuitry for affective-motivational responses to cold pain

**DOI:** 10.1101/2025.01.31.636010

**Authors:** Prannay Reddy, Takao Okuda, Mousmi Rani, Jagat Narayan Prajapati, Smriti Koul, Vijay Samineni, Arnab Barik

## Abstract

The medial thalamus is crucial for the sensory and affective-motivational responses to chronic pain. However, a mechanistic understanding of how the distinct subnuclei of the medial thalamus mediate behavioral responses to pain remains lacking. Taking advantage of intersectional viral genetics, chemogenetics, optogenetics, in-vivo imaging, and ex-vivo physiology, we reveal that the neurons in the parafascicular (PF) nuclei of the medial thalamus receive monosynaptic inputs from the lateral parabrachial nuclei (LPBN) in mice. LPBN is an essential nucleus in the ascending pain pathway, receiving projections from the dorsal horn of the spinal cord. The PF neurons downstream of LPBN (PF_post-LPBN_) are nociceptive, sensitized by peripheral neuropathy, acutely aversive, and, when activated, drive both sensory and affective-motivational responses to cold pain. In contrast, the LPBN target neurons in the intralaminar centromedian thalamus (CM_post-LPBN_), another nociceptive nucleus of the medial thalamus, are primarily involved in the affective-motivational aspects of pain. Together, we reveal that the LPBN, through two closely related thalamic nuclei, influences behavior in mice with cold hypersensitivity due to peripheral neuropathies.

## Introduction

In humans and animals, chronic pain manifests primarily in two forms. First, the threshold for tolerating somatosensory stimuli is lowered when innocuous stimuli are perceived as noxious^1^. Second, chronic pain causes long-term emotional distress, also known as the affective-motivational effects of pain^2,3^. Both of these pathological symptoms of chronic pain— hypersensitivity to sensory stimuli and adverse emotional effects are due to alterations in the neural circuits in the brain^4,5,6^. However, how the circuits in the pain-processing areas in the brain coordinate to drive these chronic pain-related pathologies remains unclear.

One of the primary targets of the spinal projection neurons that carry nociceptive information to the brain is the lateral parabrachial nucleus (LPBN)^7,8,9^. The LPBN neurons then route the noxious information across the brain^10–13^. This ascending spinoparabrachial pathway is necessary and sufficient to express affective-motivational or coping responses to pain, such as licking in mice^14,8,15^. Notably, the coping responses occur when the noxious stimuli cause damage to the integumentary system and inflict injuries, and thus are a behavioral indicator for the intensity or extent of the injury causing chronic pain^16^. This implies that the LPBN neurons can instruct pain-induced licking or coping behaviors through their downstream targets. In lesion studies in human subjects, removing parts of the medial thalamic area, particularly the centromedian-parafascicular (CM-PF) complex, alleviated chronic refractory pain^17,18^. The CM-PF complex is located within the intralaminar thalamus and comprises the caudo-medial CM and the rostro-lateral PF^19^. How the nociceptive information is transmitted to the CM and PF is still being determined. Recently, it was shown that the LPBN neurons downstream of the spinal cord primarily synapse with CM neurons^20,21^. This LPBN-CM pathway promoted affective-motivational behaviors, such as licking paws exposed to noxious stimuli. However, it remains to be tested whether the spinoparabrachial pathway communicates with the PF through the LPBN and what role this connection may play in driving the behavioral consequences of chronic pain.

Traditionally, the parafascicular thalamus (PF) is known to be involved in arousal and attention^22,23^. Recent findings indicate that the PF mediates components of motor behaviors, such as action initiation, and determines the directionality and speed of self-initiated head movements^24^. Notably, in the 1960s, seminal electrophysiological studies in cats carried out by Poggio and Mountcastle showed that the neurons in the posterior mid-thalamic areas, which include PF, are activated by noxious mechanical stimuli^25^. Meanwhile, direct stimulation of the PF neurons was found to modulate pain thresholds in humans and rats^26,27^. Here, we probed the LPBN inputs to the PF to understand how they mediate paw-licking behaviors for cold pain due to chemotherapeutic drug-induced peripheral neuropathy (CIPN). The PF neurons downstream of LPBN (PF_post-LPBN_) were activated when mice licked their paws in response to cold. PF_post-LPBN_ neurons were sufficient to accentuate the licking behavior at innocuous temperatures and were necessary for its intensity. Moreover, the PF_post-LPBN_ neurons were aversive, sufficient to cause cold pain-related negative associations, and when inhibited, impaired the ability to avoid noxious cold temperatures. Therefore, this indicates that the ascending circuitry between the LPBN and PF is vital for pain-induced protective behaviors.

### LPBN_post-spinal_ neurons innervate the PF neurons of the medial thalamus

The intralaminar thalamus (ILN), including the CM, was found to be innervated by LPBN neurons downstream of the spinal cord (LPBN_post-spinal_)^20^. Hence, we wondered if the PF neurons, the other major nuclei of the CM-PF complex, receive LPBN_post-spinal_ inputs. To that end, we labeled the LPBN_post-spinal_ neurons by stereotaxically injecting the anterograde transsynaptic AAV1-hSyn-Cre (AAVTrannsyn-Cre)^28–30^ into the lumbar segments of the spinal cord dorsal horn (SDH)^31^ and AAV5-hSyn-DIO-GFP and AAV5-hSyn-DIO-tdTomato in the left and right LPBN respectively (Figure 1A). This strategy allowed us to visualize the arborization of the LPBN neurons that receive direct synaptic inputs from the spinal cord (Figure 1B). In addition, due to the simultaneous labeling of left and right LPBN_post-spinal_ neurons with fluorescent proteins of non-overlapping fluorescent spectra, we could observe the ipsilateral-contralateral distribution of the LPBN_post-spinal_ axon terminals in the CM-PF complex (Figure 1B, Bottom panel). The LPBN_post-spinal_ neurons bilaterally innervated the CM and PF (Figure 1B, Bottom panel). Notably, the LPBN_post-spinal_ neurons express the gene for the receptor of neuropeptide Substance P, *Tacr1* (LPBN*^Tacr^*^1^), and synapses onto the CM neurons^8,20^. When we labeled the LPBN*^Tacr^*^1^ neurons with membrane-tagged GFP and mRuby fluorescent protein fused with synaptophysin in a Cre-dependent manner, we found that these neurons synapsed onto the neurons in the CM and the PF^32^ (Figure 1C). The presence of the mRuby-fused synaptophysin, along with GFP, ensured the presence of synaptic terminals and not enpassant fibers of the LPBN*^Tacr^*^1^ neurons in the CM and PF(Figure 1D).

**Figure 1.**
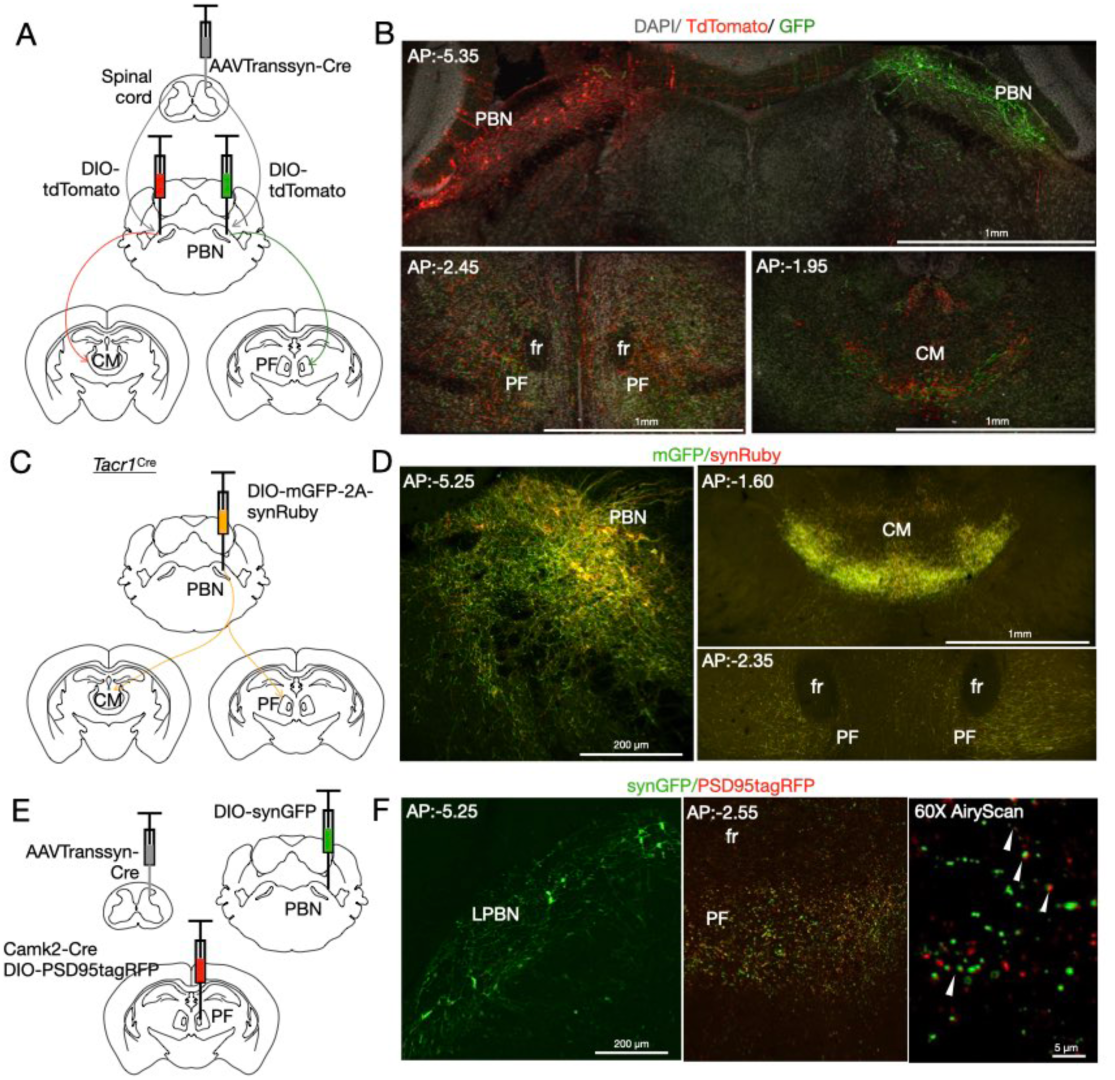
LPBN_post-spinal_ neurons synapse onto the CM and PF of the medial thalamic complex. (a) A diagram representing the viral genetic strategy used to test if the LPBN_post-spinal_ neurons innervate the PF and CM. (b) Representative confocal images of coronal sections of the LPBN (Top image), PF (Bottom left image), and CM (Bottom right image). We found that the tdTomato and GFP are expressed in the cell bodies of the LPBN_post-spinal_ neurons, while the axon terminals project bilaterally to the CM and PF. (c) A strategy combining mouse and viral genetics to test if the LPBN*^Tacr^*^1^ cells innervate the PF and CM. (d) Representative confocal images of coronal sections of the LPBN (left image), CM (top right image), and PF (bottom right image) showing the expression of the membrane-tagged GFP and synaptophysin-fused Ruby. The cell bodies and axons of the LPBN*^Tacr^*^1^ neurons were labeled with GFP, whereas the mRuby was enriched in the synaptic vesicle-laden axon terminals in the CM and PF. (e) Diagrammatic depiction of the viral genetic strategy used in wild-type mice to label the LPBN_post-spinal_ neurons with the synaptophysin-fused GFP and the post-synaptic components of the PF neurons with the PSD95-tagRFP. Notably, the PF neurons are excitatory. Hence, the PSD95-tagRFP is driven by the CamKII promoter. (f) Representative confocal images to demonstrate labeling of the LPBN_post-spinal_ neurons with the synaptophysin-fused GFP (green, left) and the closely apposed LPBN_post-spinal_ axon terminals with the PSD95-expressing dendrites of the PF (red, middle-10X and right-40X Airy scan).

The PF is rostral to the CM and is distinguished by the fiber bundles (fasciculus retroflexus) passing through it^33^. Previously, it was shown that the LPBN_post-spinal_ neurons synapse with the CM neurons; however, the anatomical data presented here provides the first evidence that LPBN_post-spinal_ neurons innervate the PF. To determine if the LPBN_post-spinal_ neurons form synapses on the PF neurons, we trannsynaptically labeled the LPBN_post-spinal_ neurons with synaptophysin-tagged GFP (synGFP) and expressed the PSD95-RFP under the CamKII promoter (labels excitatory neurons) in the PF (Figure 1E). The synGFP labeled the synaptic vesicles enriched in the axon terminals of the LPBN_post-spinal_ neurons. The PSD95-RFP was expressed in the post-synaptic densities of the PF neurons (Figure 1F). Through the AiryScan super-resolution imaging, we found that the synaptic vesicle-rich axon terminals of the LPBN_post-spinal_ neurons closely oppose the PSD95 containing post-synaptic densities of the PF neurons, indicating synapses between the two neuronal populations (Figure 1F, Right image). Collectively, our anatomical data indicate that the LPBN^post-spinal^ synapses onto neurons of CM and PF can relay nociceptive information from the SDH.

To further consolidate the findings from our anatomical studies, we performed rabies tracing from the PF neurons post-synaptic to the LPBN (PF_post-LPBN_) to test if the LPBN neurons are upstream^34,35^. We stereotaxically delivered the AAVTranssyn-Cre in the LPBN, DIO-TVA-GFP, and DIO-G in the PF of wild-type mice (Figure S1A). After two weeks, we injected the ΔG-Rabies-mCherry in the PF (Figure S1A). Anatomical characterization after one week of rabies viral injections, we found that the LPBN, rostral ventromedial medulla (RVM), and central amygdala (CeA) provide monosynaptic inputs on the PF neurons (Figure S1B). Interestingly, these nuclei are important nodes in the pain matrix, which is a map of interconnected brain nuclei that gives rise to the pain percept, and drive response to it^6^. Next, we determined the pre-synaptic inputs to the LPBN neurons projecting to the PF (LPBN_pre-PF_). We injected the retrogradely transporting AAVRetro-Cre^36^ in the PF and intersected with the DIO-TVA-GFP and DIO-G in the LPBN (Figure S1C). Stereotaxic delivery of the ΔG-Rabies-mCherry in the LPBN labeled neurons in the SDH and brain areas such as the lateral hypothalamus and PF (Figure S1D). This indicates that the LPBN_pre-PF_ neurons get direct synaptic inputs from the projection neurons in the SDH and form recurrent connections with the PF (Figure S1D, top right image).

Moreover, previous reports show that the paraventricular thalamus (PVT) and the mediodorsal parts (MD) of the thalamus are innervated directly by spinal projection neurons^37^. However, it is unclear if the CM-PF complex is innervated by ascending neurons from the spinal cord. Hence, we stereotaxically injected Cre-dependent AAV5-hSyn-DIO-tdTomato in the lumbar segments of the SDH of VGlut2*^Cre^ or Slc17a6^Cre^* mice to label the projection neurons (Figure S1E). As expected, we found axon terminals in the LPBN (Figure S1F). In addition, the SDH projection neurons terminated in the CM and PF (Figure S1G and H), indicating that direct spinal inputs, in addition to the second-order relay through the LPBN, can mediate the nociception and chronic pain pathophysiology driven by the CM-PF complex.

### LPBN neurons form functional synapses onto PF, and post-synaptic PF neurons are sensitized by oxaliplatin

Our anatomical data show that the LPBN neurons form synapses with the PF neurons (Figure 1). Hence, next, we determined if the LPBN and CF are functionally connected. Further, we explored if the electrophysiological properties of the PF_post-LPBN_ neurons are altered in CIPN mice. We hypothesized that the synaptic inputs, specifically from the LPBN neurons onto the PF_post-LPBN_ neurons is strengthened post-intraplantar oxaliplatin administration. In that case, the changes in the PF_post-LPBN_ inputs can explain the enhanced licking observed in mice with CIPN^15^. To that end, we injected AAVTranssyn-Cre in the LPBN of the Ai213 transgenic strain^38^ (Figure 2A). In Ai213 mice, Cre recombinase conditionally expresses GFP in the cell type of choice. Thus, as expected, we observed GFP-expressing cells in the PF, indicating that a subpopulation of PF neurons receives inputs from the LPBN (Figure 2B). Next, we performed voltage-clamp intracellular recordings from the PF_post-LPBN_ neurons in the Ai213 mice. We found that the spontaneous firing properties of the GFP expressing PF_post-LPBN_ neurons are accentuated post-injection of intraplantar oxaliplatin (Figure 2D). The amplitude is increased, and the inter-event interval of the mEPSCs is decreased in the PF_post-LPBN_ neurons of mice with CIPN (Figure 2E). This indicates the increase in the strength of the synaptic inputs onto the PF_post-LPBN_ neurons in mice with CIPN. However, the transsynaptic labeling strategy results do not confirm the specific involvement of the LPBN inputs in the increased pre-synaptic inputs to the PF neurons due to intraplantar oxaliplatin administration. To address this, we expressed the light-sensitive optogenetic actuator channelrhodopsin-2 (ChR2)^39^ in the LPBN neurons. We tested if the light-evoked EPSCs in the PF_post-LPBN_ are altered in mice with neuropathic cold allodynia (Figure 2F). We found that the amplitudes of the light-evoked EPSCs are increased in mice with CIPN (Figure 2G). This observation reaffirms the functional connectivity between LPBN and PF and demonstrates that the synaptic strength between the PBN and PF changes in mice with peripheral neuropathy.

**Figure 2.**
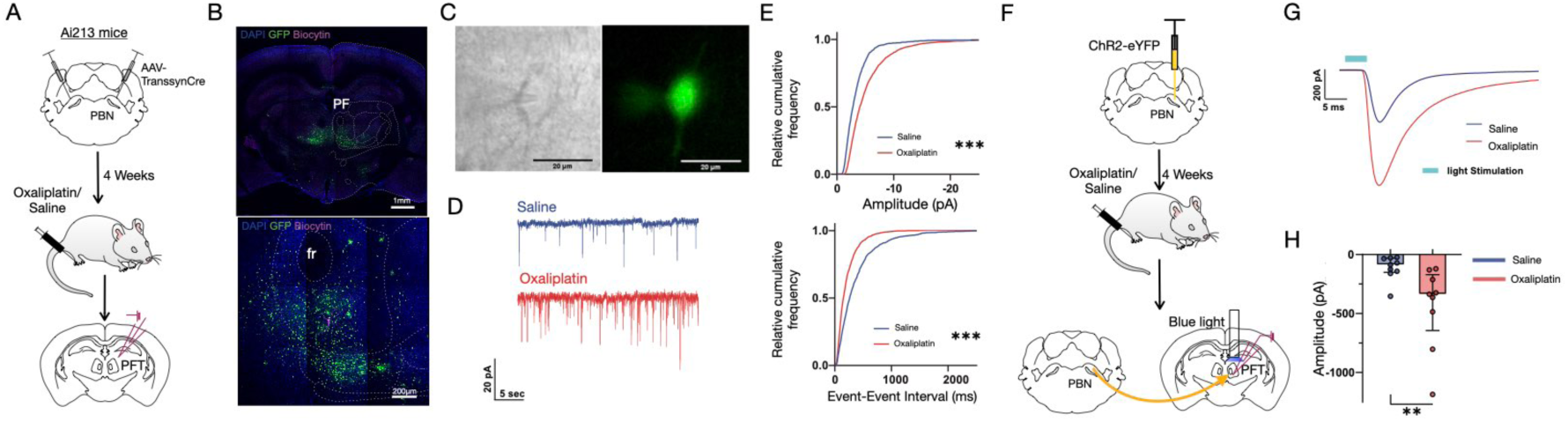
Functional synaptic connectivity between the LPBN and PF. (a) A schematic of the time course of the patch clamp recordings from the PF_post-LPBN_ neurons. AAVTranssyn-Cre was injected in Ai213 mice in the LPBN, and after 4 weeks, intraplantar injections of oxaliplatin were performed in the hind-paw to induce CIPN. Brains were then harvested from the mice of ex-vivo slice electrophysiology.(b) A confocal image of a coronal section of the PF_post-LPBN_ cells expressing GFP (Top image) used for recordings and filled with Biocytin (Bottom image). (c) Representative images of whole-cell patch clamp of PF_post-LPBN_ neurons. The left image shows the IR-DIC image and the right image shows the GFP fluorescent image. (d) Representative traces of voltage-clamp recordings at −70mV for saline (Top trace in blue) and oxaliplatin (Bottom trace in red) injected mice. (e) Cumulative relative frequency plot of mEPSC amplitude (Top plot) and mEPSC event-to-event interval time (Bottom plot) for all events recorded. Red trace: total 2608 events from 3 neurons in 2 Oxaliplatin-treated mice. Blue trace: 1494 events from 3 neurons in 2 Saline-treated mice. ***p<0.0001, two-sample Kolmogorov–Smirnov test. (f) A schematic of the time course of the light-evoked EPSC recording in the PF. AAV-hSyn-ChR2-eYFP virus was injected into the LPBN 4 weeks before the patch-clamp recording, and then the mice were injected with oxaliplatin or saline. (g) Representative traces of light-evoked EPSC from each group. The blue sample trace is from the saline-treated mouse, and the red trace is from the Oxaliplatin-treated mouse. Each trace is the average trace of 10 stimuli. (h) A plot of the amplitude of light-evoked EPSCs for each group. Each dot represents each recorded neuron (9 neurons from 3 Saline-treated mice and 9 neurons from 2 Oxaliplatin-treated mice). Each filled bar indicates the median, and each error bar indicates the interquartile range of each group. **p = 0.008, Mann-Whitney test.

### Activation of LPBN terminals in PF and CM regulates nocifensive behaviors

Recently, we have shown that the LPBN_post-spinal_ neurons mediate nocifensive behaviors such as licking in response to cold pain^15^. In mice with CIPN, chemogenetic activation of the LPBN_post-spinal_ neurons was sufficient to promote licking behavior in response to cold^40^. We found that the frequency of licks and the length of time for which the lick bouts lasted were increased in mice when the LPBN_post-spinal_ (Figure S2A, B) or the entire LPBN neurons were chemogenetically activated (Figure S3). We interpreted the number of licks during the assay as the frequency with which the pain sensation crossed the threshold to engage in attentive behaviors such as licking, whereas the length of time for which the licking lasted or bouts as the intensity with which the perception of pain occurred. The latency to lick determined the thresholds for cold allodynia in mice with CIPN.

Here, we tested if activating the LPBN terminals in the CM or PF affects the licking behavior in mice with CIPN on the cold-plate test (Figure 3A, B). We expressed Channelrhodopsin-2 (ChR2) in the LPBN neurons and optogenetically stimulated the axon terminals in the CM and PF. We injected AAV-DIO-ChR2-mCherry in the right LPBN of *Slc17a6^Cre^* mice and implanted optic fiber on either PF or CM to excite the LPBN axon terminals in these two anatomically distinct nuclei of the medial thalamus (Figure 3B, C). We found that activating the LPBN terminals in the PF increased the frequency and decreased the latency of licks without affecting the bouts on the gradient (24°C-0°C over 5 minutes) and static (12°C over 5 minutes) cold-plate tests in mice with CIPN (Figure 3D-G). On the other hand, activating the LPBN terminals in the CM increased the frequency (Figure 3H-L). The bout lengths and latency were unaffected. Mice without CIPN did not lick their paws on the cold-plate tests; hence, control experiments without intraplantar oxaliplatin were excluded^40^. Thus, the CM and PF neurons receive nociceptive input from the LPBN. Yet, the two inputs impart distinct (sensory) and compensatory (affective-motivational) effects on pain-induced licking behavior.

**Figure 3.**
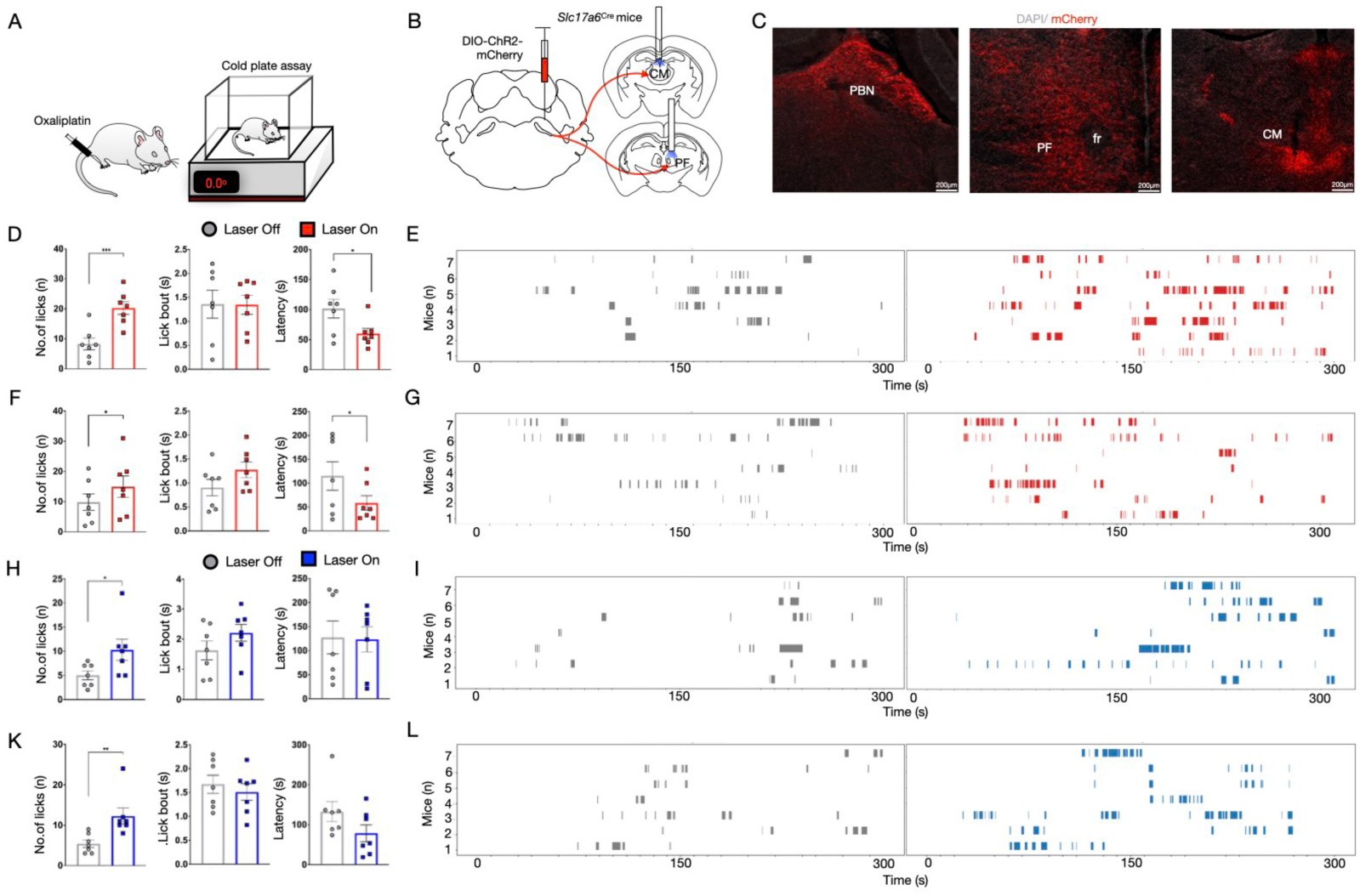
Optogenetic activation of LPBN terminals in the CM and PF accentuates coping behaviors. (a) A schematic of the intraplantar injection of oxaliplatin (CIPN) or saline (control) and the thermal plate assay to test for nociceptive thresholds and nocifensive behaviors. (b) A diagrammatic representation of the strategy used to stimulate the LPBN projections in the PF or CM optogenetically. pAAV-Ef1a-DIO hChR2(C128S/D156A)-mCherry was injected in the LPBN in *Slc17a6^Cre^* mice, and fiber was stereotaxically implanted over the PF or CM. (c) Representative confocal images of the coronal sections of the LPBN (left image), PF (center image), and CM (right image) showing expression of mCherry tagged ChR2 (red). (d) Quantification of lick frequency (8.28±1.96 compared to 20.29±2.10, respectively; t-test, \*\*\**P*=0.0008), the length of lick bouts (1.36±0.29 compared to 1.35±0.20, respectively) and latency to lick (101.71±15.65 compared to 60.33±8.45, respectively; t-test, \**P*=0.02) on the gradient cold plate assay (24 to 0°C over 5mins) with laser on to excite the LPBN terminals in PF or laser off as the controls (n=7). (e) Raster plots showing the lick occurrences across a single session of gradient cold plate assay with laser off (left plots) or laser on to excite the LPBN terminals in PF (right plots) (n=7). (f) Quantification of lick frequency (9.86±3.55 compared to 15.00±2.70, respectively; t-test, \**P*=0.03), the length of lick bouts (0.90±0.17 compared to 1.27±0.16, respectively) and latency to first lick (115.01±29.96 compared to 58.76±15.17, respectively; t-test, \**P*=0.03) on the static cold plate assay (12°C over 5mins) with laser on to excite the LPBN terminals in PF or laser off (n=7). (g) Raster plots for licks across a single session of static cold plate assay with laser off (controls, left plots) or laser on to excite the LPBN terminals in PF (right plots) (n=7). (h) Quantification of lick frequency (5.00±0.90 compared to 10.29±2.18, respectively; t-test, \**P*=0.03), the length of lick bouts (1.62±0.32 compared to 2.21±0.28, respectively) and latency to lick (127.56±34.24 compared to 123.54±26.38, respectively) on the gradient cold plate assay (24 to 0°C over 5mins) with laser on to excite the LPBN terminals in CM or laser off (n=7). (i) Raster plots demonstrating licks across a single session of gradient cold plate assay with laser off (controls, left plots) or laser on to excite the LPBN terminals in CM (right plots) (n=7). (j) Quantification of lick frequency (5.43±0.90 compared to 12.29±2.01, respectively; t-test, \*\**P*=0.007), the length of lick bouts (1.67±0.19 compared to 1.51±0.17, respectively) and latency to lick (132.33±25.07 compared to 78.23±21.06 respectively) on the static cold plate assay (12°C over 5mins) with laser on to excite the LPBN terminals in CM or laser off (n=7). (k) Raster plots for licking across a single session of static cold plate assay with laser off (left plots) or laser on to excite the LPBN terminals in CM (right plots) (n=7).

Next, we explored the projections of the PF and CM neurons downstream of the LPBN. We hypothesized that the output patterns of the PF_post-LPBN_ and CM_post-LPBN_ may explain the differences in cold pain-induced nocifensive behaviors. To that end, we injected AAVTranssyn-Cre in the LPBN and DIO-tdTomato in the CM and DIO-GFP in the PF of wild-type mice (Figure S4A). As a result, we found that the CM_post-LPBN_ and PF_post-LPBN_ neuronal cell bodies were labeled by GFP and tdTomato, respectively (Figure S4B). We observed the CM and PF axon terminals in the LPBN, RVM, and striatum (Figure S4C). The CM_post-LPBN_ neurons targeted the medial PBN, whereas the PF_post-LPBN_ neurons sent projections to the LPBN (Figure S4C). The PF_post-LPBN_ axons were found to innervate the RVM (Figure S4C). In the striatum, the CM_post-LPBN_ neurons terminate in the dorsal and medial parts, whereas the PF_post-LPBN_ neurons project to the ventral and lateral coordinates (Figure S4C). The CM-PF complex is known to have dense striatal projections^22,41^. Here, we show that the nociceptive neurons downstream of LPBN in CM or PF have distinct projections to the striatum and, hence, can play independent roles in nocifensive behaviors.

### PF_post-LPBN_ neurons are nociceptive and tuned to licking behaviors

Diverse sensory stimuli, including high-threshold somatosensory stimuli, are known to stimulate the PF neurons. These studies were done in rats and primates^42–44^. However, it is not known if mice PF neurons are nociceptive. To that end, we carried out microendoscopic calcium imaging of the PF neurons in lightly anesthetized (0.25% isofluorane) mice to visualize the calcium dynamics of PF neurons as they were pinched at the tail (Figures 4A and B). We injected the Cre-dependent AAV encoding the genetically encoded calcium indicator, GCaMP6s^45^, in the PF of *Slc17a6^Cre^* mice and implanted a GRIN lens over the PF to visualize the neurons through a head-mounted microendoscope^45^ (Figure 4C). Pinching the tail with forceps instead of holding it (touch) evoked calcium transients, indicating PF activation by noxious mechanical somatosensory stimuli and not by gentle mechanical stimuli (Figure 4D). Next, we sought to correlate the neural activity of the PF_post-LPBN_ neurons to the attentive behaviors elicited by cold stimuli in mice with CIPN. To test if the cold-pain induced licking engages the PF_post-LPBN_ neurons, we transsynaptically labeled the neurons with GCaMP6s (Figure 4E). Additionally, we co-expressed the non-Cre dependent jRGECO1a^46^. in the PF of the same mice to label neurons irrespective of the connectivity with LPBN (Figure 4E and F). We simultananeously imaged the red and green calcium indicators expressed in the PF neurons in behaving mice as they underwent experiences of cold allodynia with the fiber photometry technique^47^. The GCaMP6s expressing PF^post-LPBN^ neurons were found to be precisely tuned to the paw-licking caused by cold allodynia (Figures 4I and L). In contrast, the jRGECO1a expressing PF neurons were engaged by movement irrespective of licking (Figure 4J and M). Thus, we found that during licking, PF^post-LPBN^ neurons were activated, and not during general exploration of the home cage. Meanwhile, the PF neurons which do not receive synaptic inputs from the LPBN were tuned to movements that included exploration and licking (Figure 4J and M). Interestingly, we found that the PF neurons do not respond to shakes, indicating that these neurons may specifically mediate supraspinal behaviors such as licking and not shakes driven by spinal reflexes (Figure 4I, middle plots). We also tested if the PF_post-LPBN_ neurons respond to mechanical stimuli. Our recordings show that pinching the tail with forceps compared to gentle mechanical stimuli provided by grabbing with the forceps engaged the PF_post-LPBN_ neurons (Figure S5A-D). Additionally, we found that the PF neurons preferentially respond to contralateral noxious somatosensory stimuli (Figure S5E-H). Thus, our data demonstrate that the PF_post-LPBN_ neurons are engaged by cold allodynia and specifically activated when mice licked their paws in response to pain.

**Figure 4.**
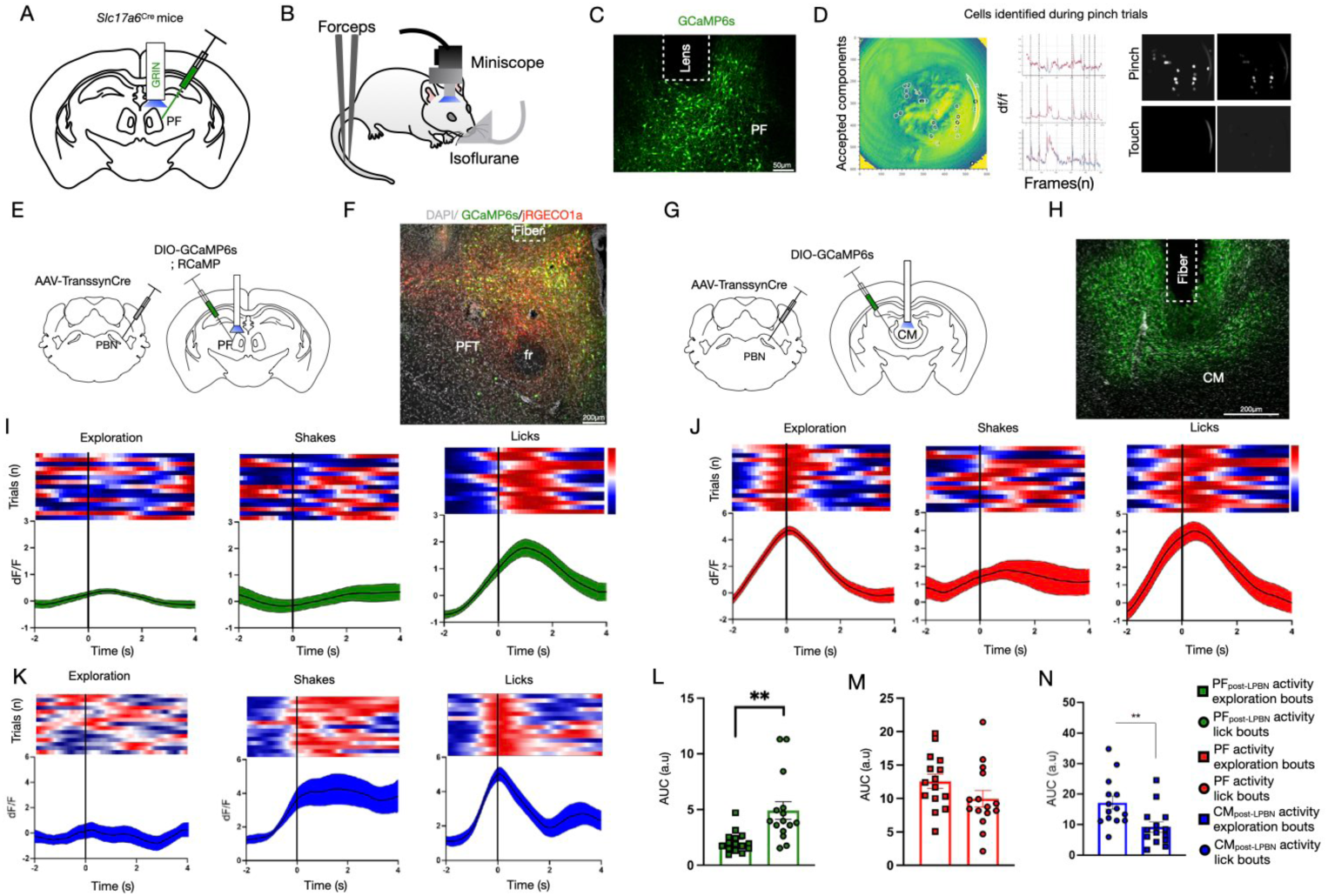
PF neurons are activated by cold allodynia and paw-licking. (a) Diagram showing stereotaxic delivery of AAV5-hSyn-DIO-GCaMP6s and subsequent GRIN lens implantation in the PF of *Slc17a6^Cre^*mice. (b) Miniscope recordings were done from the PF through the GRIN lens in mice under isofluorane anesthesia. Tips of blunt forceps were used to hold the tail to deliver a gentle touch and pressed against with force to deliver the pinch stimuli. (c) Image demonstrating GCaMP6s expression on the PF of the *Slc17a6*^Cre^ mice. The lens track suggests a successful implant over the PF. (d) Field of view through the GRIN lens with selected ROIs (left), the activity of multiple selected cells show activity in response to forceps pinches (center), and maximum intensity projections of activity of the PF neurons in response to pinching (right, top), and touch (right, bottom). (e) Strategy used to record from the PF and PF_post-LPBN_ neurons simultaneously. AAVTranssyn-Cre was injected in the LPBN, and AAV5-hSyn-DIO-GcaMP6s and AAV1-CAG-jRGECO1a were injected into the PF with a fiber. (f) A representative confocal image of a coronal section of the PF showing expression of both GCaMP5s (green) and jRGECO1a (red). (g) A diagrammatic representation of the strategy used to record from the CM. AAV5-hSyn-DIO-GcaMP6s was injected in the CM (what mice), and a fiber optic cannulae was surgically inserted on the injection site. (h) A representative confocal image of a coronal section of the CM showing expression of GCaMP6s (green). (i) Heatmaps of single trials and cumulative traces of the GCaMP6s activity in the PF_post-LPBN_ cells during exploration bouts, paw shakes, and lick bouts in mice with intraplantar injection of oxaliplatin. (n=2) (j) Heatmaps of single trials and cumulative traces of the jRGECO1a activity in the PF during exploration bouts, paw shakes, and lick bouts in mice with intraplantar oxaliplatin injection. (n=2) (k) Heatmaps of single trials and cumulative traces of the GCaMP6s activity in the CM during exploration bouts, paw shakes, and lick bouts in mice with intraplantar oxaliplatin injection. (n=4) (l) Quantification comparing the average GCaMP6s activity (AUC) during exploration bouts vs lick bouts (2.07±0.24 compared to 4.91±0.79, respectively; t-test, \*\**P*=0.002) in the PF_post-LPBN_ cells (n=2). (m) Quantification comparing the average jRGECO1a activity (AUC) during exploration bouts vs lick bouts (12.57±1.06 compared to 9.96±1.23, respectively) in the PF (n=2). (n) Comparison between the GCaMP6s activity in the CM during exploration bouts and lick bouts. Calculated from the AUC of the cumulative traces. (17.14±1.98 compared to 9.20±1.57, respectively; t-test, \*\**P*=0.007) (n=4)

Next, we carried out fiber photometry recordings from the CM neurons downstream of the LPBN (CM_post-LPBN_) (Figure 4G). We trannsynaptically labeled the CM_post-LPBN_ neurons with GCaMP6s (Figure 4G, H), and when we recorded the mice with CIPN on the cold plate test, we found that the CM_post-LPBN_ neurons responded to both shakes and licks (Figure 4K). Remarkably, the activity of the CM_post-LPBN_ neurons persisted after the mice shook their paws (Figure 4K, center plots). In contrast, the activity of the CM_post-LPBN_ neurons rose when the mice licked their paws and dropped after the lick bout was over (Figure 4K, right plots). Similar to the recordings obtained from the PF_post-LPBN_, the CM_post-LPBN_ neurons did not respond vigorously to exploratory behavior (Figure 4K, left plots, N). Thus, cold pain engages the CM_post-LPBN_ neurons during both the reflexive and coping nocifensive behaviors.

### PF_post-LPBN_ neurons mediate somatosensory and affective-motivational responses to cold pain

Next, we tested if acute activation of the PF_post-LPBN_ neurons will affect the licking behavior due to cold pain, and compared the roles played by the PF_post-LPBN_ and CM_post-LPBN_ neurons. To stimulate the PF_post-LPBN_ and CM_post-LPBN_ neurons transiently and specifically, we expressed the ChR2 with the Cre-dependent transsynaptic strategy described previously (Figure 5A). We activated the PF_post-LPBN_ and CM_post-LPBN_ neurons by shining a blue light through optogenetic cannulae placed above the PF and CM, respectively (Figure 5B and C). In the optogenetic activation experiments, we found that when the PF_post-LPBN_ neurons were stimulated and mice were placed on the gradient cold-plate test assay, the mice with CIPN demonstrated a shorter latency to lick (Figure 5D, right plot), with increased frequency and unchanged licking bouts (Figure 5D, left plot). In contrast, when we activated the CM_post-LPBN_ neurons, the lick bouts were increased, keeping the frequency and latency unaltered (Figure 5E). Thus, the PF_post-LPBN_ neurons mediate the somatosensory and affective-motivational components of cold allodynia induced nocifensive behaviors. Meanwhile, the CM_post-LPBN_ neurons are involved solely in affective-motivational aspects. Since we interpreted the length of lick-bouts as the intensity of felt pain, the CM_post-LPBN_ neurons are likely involved in expressing the vigor with which coping responses are generated. We further buttressed our findings by testing the effects of chemogenetic activation of the PF_post-LPBN_ and CM_post-LPBN_ neurons on cold pain-driven defensive behaviors. To that end, we expressed the excitatory chemogenetic actuator^48,49^, hM3Dq in the CM_post-LPBN_ and PF_post-LPBN_ neurons by injecting AAVTrannsyn-Cre in the LPBN and DIO-hM3Dq in the CM or PF of wild-type mice (Figure 5F, G). Upon i.p. administration of the high-affinity hM3Dq ligand, deschloroclozapine (DCZ)^50^ in the hM3Dq expressing CIPN-mice, we found that the activation of the PF_post-LPBN_ neurons increased the frequency of licking and reduced the latency in response to cold stimuli (Figure 5H). Whereas chemogenetic stimulation of the CM_post-LPBN_ increased the bout lengths of licking. Both optogenetic and chemogenetic strategies revealed how the PFpost-LPBN and CMpost-LPBN can regulate pain-induced coping behaviors such as paw licking in mice.

**Figure 5.**
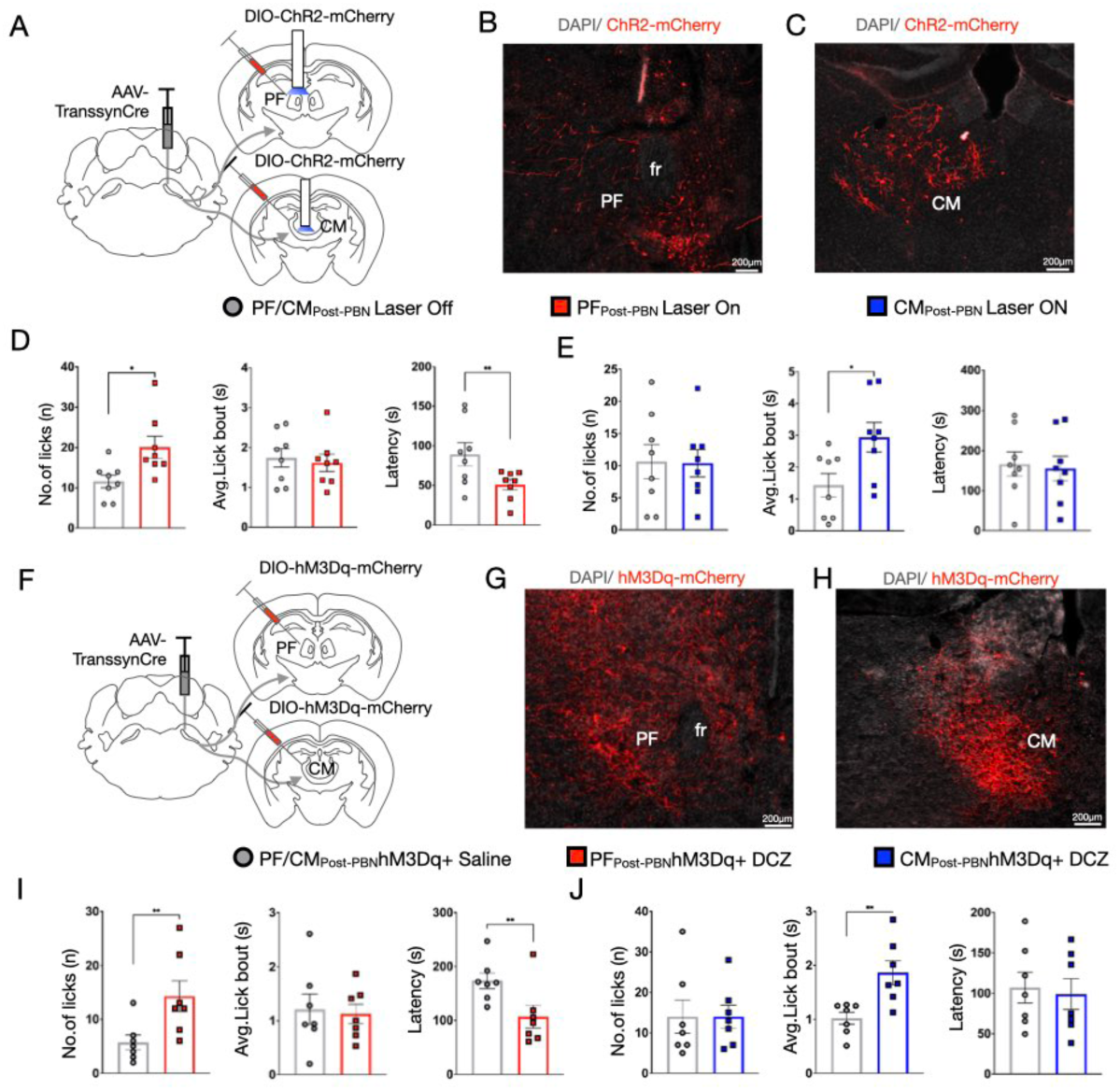
Chemogenetic activation of the PF_post-LPBN_ and CM_post-LPBN_ neurons affects licking to cold pain in mice with CIPN. (a) Strategy to selectively stimulate the PF or CM neurons post-synaptic to the LPBN. AAVTrannsyn-Cre was injected in the LPBN and pAAV-Ef1a-DIO hChR2(C128S/D156A)-mCherry with a fiber in either PF or CM. (b) Representative confocal image of a coronal section of the PF and (c) CM showing expression of ChR2 with the promoter mCherry(red). (d) Quantification of lick frequency (11.63±1.55 compared to 20.13±2.68, respectively; t-test, \**P*=0.01), length of lick bouts (1.74±0.23 compared to 1.61±0.22, respectively) and latency to lick (89.38±14.71 compared to 50.90±6.49, respectively; t-test, \*\**P*=0.009) on the gradient cold plate assay (24 to 0°C over 5mins) with laser on to excite the PF_post-LPBN_ cells or laser off as controls (n=8). (e) Quantification of lick the frequency (10.63±2.67 compared to 10.38±2.12, respectively), lengths of the lick bouts (1.43±0.36 compared to 2.94±0.46, respectively; t-test, \**P*=0.02) and latency to lick (166.36±30.72 compared to 155.21±31.14, respectively) on the gradient cold plate assay (24 to 0°C over 5mins) with laser on to excite the CM_post-LPBN_ cells or laser off as controls. (n=8) (f) Strategy to chemogenetically stimulate the PF or CM targets of LPBN. AAVTranssyn-Cre was injected in the LPBN and AAV5-hSyn-hM3D(Gq)-mCherry in either PF or CM. (g) A representative confocal image of a coronal section of the PF and (h) CM showing expression of hM3Dq fused with mCherry(red). (i) Quantification of lick frequency (5.71±1.38 compared to 14.29±2.85, respectively; t-test, \*\**P*=0.003), length of the bouts (1.21±0.28 compared to 1.13±0.18, respectively) and latency to lick (173.23±14.86 compared to 106.44±20.85, respectively; t-test, \*\**P*=0.006) on the gradient cold plate assay (24 to 0°C over 5mins) after i.p injection of saline or DCZ (n=7). (j) Quantification of lick frequency (14.00±4.09 compared to 14.00±2.83, respectively), lengths of the lick bouts (1.02±0.11 compared to 1.86±0.22, respectively), and latency to lick (107.00±19.13 compared to 99.06±18.92, respectively) on the gradient cold plate assay (24 to 0°C over 5mins) with an i.p injection of saline or an i.p injection of DCZ in mice with hM3Dq in the CM_post-LPBN_ neurons (n=7).

### Activation of the PF_post-LPBN_ neurons drives acute and learned aversion

Next, we explored if the PF_post-LPBN_ neurons play a role in acute and learned aversion. We performed the real-time place aversion (RTPA) test on mice expressing ChR2 in the PF_post-LPBN_ and CM_post-LPBN_ neurons (Figure 6A). In the RTPA test, carried out with optogenetic stimulation paradigms (Figure 6B), the neuronal population of interest is stimulated when mice enter a designated chamber in a two-chamber apparatus of identical design. It is then determined if the optogenetic stimulation of target neurons evokes escape or promotes aversion of the paired chamber by calculating the time spent. We found that optogenetic activation of the PF_post-LPBN_ and not CM_post-LPBN_ neurons promoted RTPA from the paired chamber (Figure 6F). This finding indicates that the PF_post-LPBN_ neurons are involved in acutely aversive behaviors. The conditioned place aversion test (CPA) for mice with Cre recombinase-dependent chemogenetic activation strategies allows for testing if a neural population is sufficient in causing learned aversion. Thus, next, we tested if the PF_post-LPBN_ and CM_post-LPBN_ neurons can drive mice to learn to avoid a paired chamber on the CPA test. We paired mice expressing hM3Dq in the PF_post-LPBN_ and CM_post-LPBN_ neurons with a striped chamber with saline (control) or the DREADDs ligand DCZ (Figure 7B). We found that, in comparison with the saline-treated mice, when the mice with hM3Dq in the PF_post-LPBN_ or CM_post-LPBN_ neurons were treated with DCZ in the striped chamber, the mice spent less time in the striped chamber (Figure 7E and H)—indicating that the chemogenetic activation of both the PF_post-LPBN_ and CM_post-LPBN_ neurons is sufficient for forming a negative association with a paired chamber in the CPA test. In sum, we conclude that the PF_post-LPBN_ are sufficient to drive acute and learned aversion, whereas the CM_post-LPBN_ neurons mediate learned aversive behaviors and not the acute ones.

**Figure 6.**
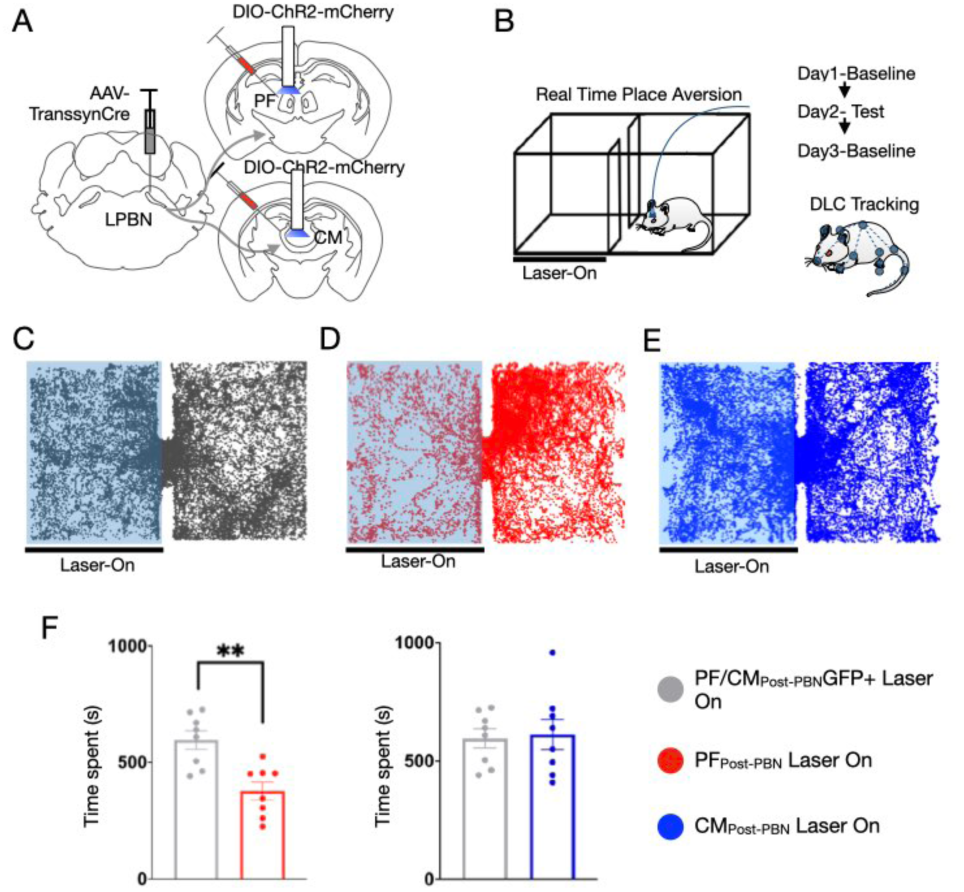
Chemogenetic activation of the PF_post-LPBN_ and CM_post-LPBN_ neurons causes CPA. (a) Strategy to optogenetically stimulate the PF or CM neurons in wild-type mice. AAVTranssyn-Cre was injected in the LPBN and pAAV-Ef1a-DIO hChR2(C128S/D156A)-mCherry with a fiber in either PF or CM. (b) Schematic representation of the experimental paradigm and setup for the RTPA assay. (c) Representative traces tracking the movement of the mice in the RTPA box for mice injected with GFP in PF_post-LPBN_ or CM_post-LPBN_ cells (sham), (d) ChR2 in PF_post-LPBN_ cells, and (e) ChR2 in CM_post-LPBN_ cells. (f) Quantification comparing the time spent on the laser ON side of the RTPA box for GFP (controls) vs. ChR2 expressing PF_post-LPBN_ cells (596.00±39.84 compared to 377.00±38.21, respectively; t-test, \*\**P*=0.001) (Left plot) and GFP vs. ChR2 expressing CM_post-LPBN_ neurons (596.00±39.84 compared to 613.25±63.56, respectively) (Right plot).(n=8)

**Figure 7.**
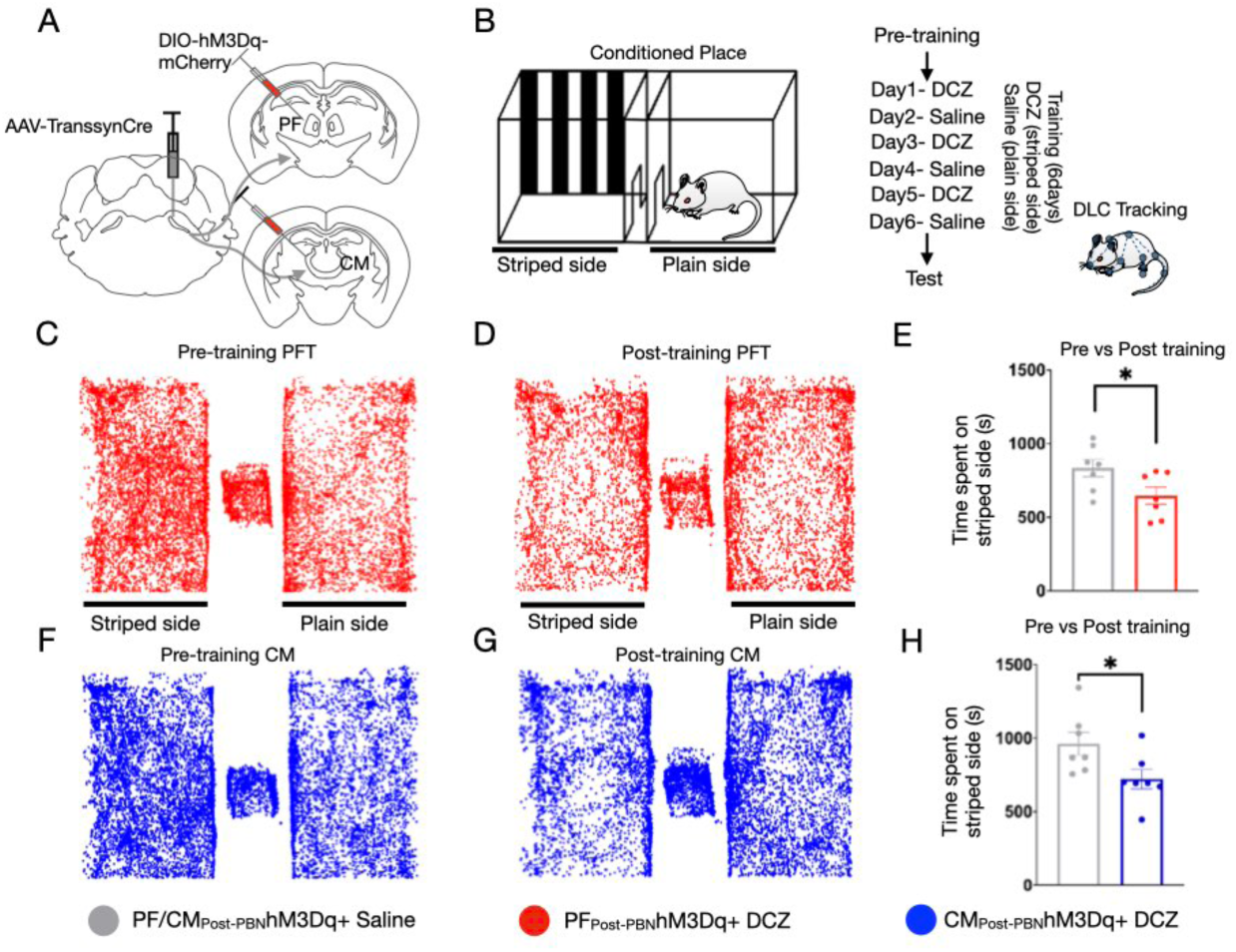
Optogenetic activation of the PF_post-LPBN_ neurons is acutely aversive. (a) Representation of the strategy used to stimulate the PF or CM neurons chemogenetically. AAVTrannsyn-Cre was injected in the LPBN and AAV5-hSyn-hM3D(Gq)-mCherry in either PF or CM. (b) Schematic explaining the experimental setup and paradigm of the CPA assay. (c) A representative trace of mouse tracking in the CPA assay showing the time spent in the striped and plain side before training and (d) after training in mice expressing hM3Dq in the PF_post-LPBN_ neurons. (e) Quantification of the time spent on the side of the CPA box by mice that received an i.p injection of DCZ pre-vs post-training (test) (833.86±59.10 compared to 646.29±57.75, respectively; t-test, \**P*=0.04) in mice with hM3Dq in the PF_post-LPBN_ neurons (n=7). (f) A representative trace of mouse tracking in the CPA assay showing the time spent in the striped and plain side before training and (g) after training in mice with hM3Dq in the CM_post-LPBN_ neurons. (h) Quantification comparing the time spent on the side of the CPA box mice received an i.p injection of DCZ pre- vs post-training (test) (962.29±78.07 compared to 695.00±65.28, respectively; t-test, \**P*=0.04) in mice expressing hM3Dq in CM_post-LPBN_ neurons(n=7).

### PF_post-LPBN_ neurons are required for the expression of coping responses to pain

We found that the activation of PF_post-LPBN_ and CM_post-LPBN_ neurons differentially affects the frequency, latency, and bout lengths of licks induced by cold stimuli in mice with CIPN (Figure 5). Hence, we asked how the inhibition of these neurons affects the coping responses to cold pain. We expressed the inhibitory DREADDs^48^ (hM4Di) in the PF_post-LPBN_ and CM_post-LPBN_ neurons to enable silencing upon i.p. administration of DCZ (Figure 8A). Saline administration in the same mice served as controls. Upon silencing both the PF_post-LPBN_ and CM_post-LPBN_ neural populations, we found that the length of licking bouts on the cold plate was reduced (Figure 8C, center plot). Meanwhile, the number and latency to lick were unaltered (Figure 8C, left and right plots). We confirmed this finding by silencing the presynaptic inputs (reducing the excitability) to the PF_post-LPBN_ and CM_post-LPBN_ neurons by selectively expressing Kir2.1 channels^51^. Mice expressing GFP in the PF_post-LPBN_ and CM_post-LPBN_ were used as controls. Compared to the GFP controls, mice expressing Kir2.1 in the PF_post-LPBN_ and CM_post-LPBN_ displayed reduced bout lengths on the gradient-cold plate test after developing CIPN (Figure 8F, center plot). Together, we observed that acute (hM4Di) or chronic (Kir2.1) inhibition of the PF_post-LPBN_ and CM_post-LPBN_ neurons reduces the lick-bout lengths in mice with cold-allodynia, indicating their involvement in experiencing pain intensity. Next, we tested if the chemogenetic inhibition of the PF_post-LPBN_ neurons would be altered in the temperature-plate preference (TPP) test when mice with CIPN could choose between two thermal plates set at 4°C and 24^o.^C (Figure 8G). We previously observed that PF_post-LPBN_ neurons, when activated, reduced the latency to lick to cold stimuli and increased the frequency of licks (Figure 5D). Hence, we hypothesized that inhibiting the PF_post-LPBN_ neurons may alter the place preference in CIPN mice if they can choose between two plates set at innocuous (24°C) and noxious (4°C) cold temperatures on the TPP test. Indeed, neuropathic mice with i.p. DCZ, in comparison with the saline-injected controls, spent more time on the 4°C plate (Figure 8H). This indicates that the transient inactivation of the CM_post-LPBN_ neurons is sufficient to partially reverse the cold hypersensitivity due to oxaliplatin-induced CIPN.

**Figure 8.**
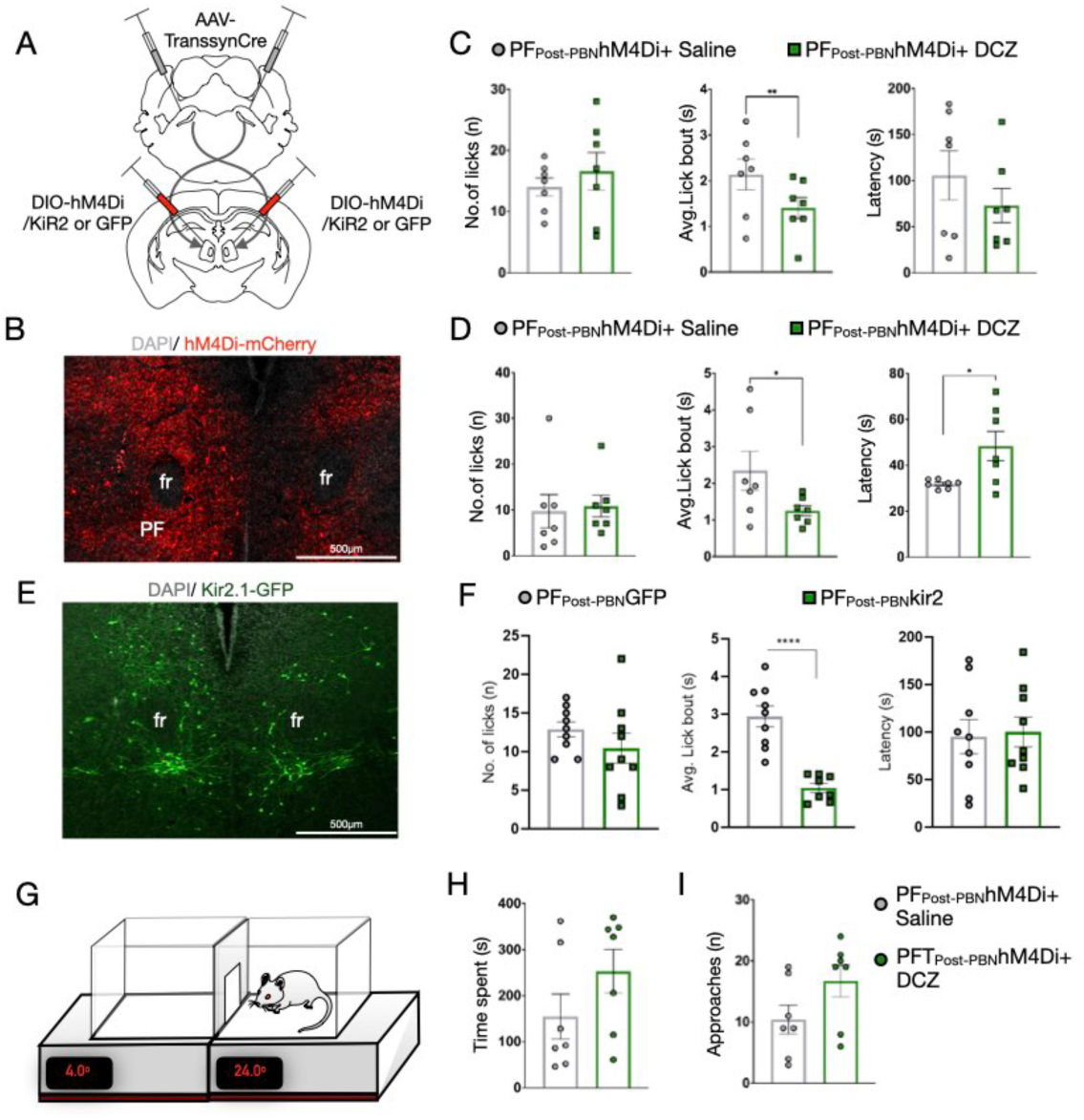
Inhibition of the PF_post-LPBN_ neurons affects the length of lick bouts and aversion due to cold hypersensitivity. (a) Diagram representing the strategy used for the bilateral inhibition of PF_post-PBN_ neurons. AAVTranssyn-Cre was injected in the LPBN of both the hemispheres, and AAV5-hsyn-DIO-hM4D(GI)-mCherry or AAV-EF1a-DIO-Kir2.1-P2A-EGFP-WPRE-hGH bilaterally in the PF. (b) A confocal image of a coronal section of the PF shows the expression of the mCherry fused hM4Di (red). (c) Quantification of lick frequency (14.00±1.48 compared to 16.57±3.09, respectively), length of the lick bouts (2.14±0.34 compared to 1.41±0.23, respectively; t-test, \*\**P*=0.007) and the latency to lick (105.71±26.58 compared to 72.93±18.44, respectively) on the gradient cold plate assay (24 to 0°C over 5mins) with an i.p injection of saline or an i.p injection of DCZ to inhibit the PF_post-LPBN_ cells. (n=7) (d) Quantification of the lick frequency (9.71±3.64 compared to 10.86±2.39, respectively), length of the lick bouts (1.26±0.14 compared to 2.34±0.53, respectively; t-test, \*\**P*=0.047) and latency to lick (31.87±0.72 compared to 48.39±6.38, respectively; t-test, \**P*=0.03) on the static cold plate assay (24 to 0°C over 5mins) with an i.p injection of saline or an i.p injection of DCZ. (n=7) (e) A representative confocal image of a coronal section of the PF showing the expression of KiR with a fluorescent tag GFP (green). (f) Quantification of lick frequency, length of the lick bouts (t-test, \*\*\*\**P*=0.000), and latency to lick on the gradient cold plate assay (24 to 0°C over 5mins). (n=9) (g) A Temperature Place Preference (TPP) assay thermal plate schematic with the mouse placed on the 24°C (innocuous) surface. (h) Comparison of the time spent on the 4°C side of the TPP test (154.57±48.97 compared to 253.14±47.49, respectively) and (i) the number of approaches to the entrance of the 4°C side (10.43±2.35 compared to 16.71±2.60, respectively) for an i.p injection of saline (controls) vs an i.p injection of DCZ to inhibit the PF_post-LPBN_ cells. (n=7)

## Discussion

This study explored how neural connectivity between the LPBN and PF of the CM-PF complex mediates the behavioral responses to cold pain in mice with CIPN. We particularly focused on the licking behavior, as it is one of the innate coping mechanisms through which mice manage pain. We provide the first evidence that a subset of PF neurons, receiving synaptic inputs from the LPBN, are specifically tuned to noxious somatosensory stimuli and licks. The LPBN-PF neural pathway is potentiated in animals with peripheral neuropathy, and when artificially manipulated, it affects cold pain-induced licking behavior. Finally, this circuitry is sufficient for acute and learned aversion. Furthermore, we delineated a novel brainstem-thalamic circuitry that plays a vital role in expressing affective-motivational behaviors in animals with chronic pain.

Traditionally, it is thought that the spinal projections to the LPBN synapse onto the central amygdala (CeA) projecting neurons^52^. However, recently, it was shown that the LPBN neurons receiving spinal inputs project to the intralaminar thalamus, which includes the CM^20^. The CeA projecting LPBN neurons express *Calca* and *Tac1*^52,10^, whereas the CM-PF projection neurons express *Tacr1*^20,8^. Noxious somatosensory stimuli activate all of the three LPBN neural populations. Thus, the LPBN-CeA pathway is potentially indirectly activated and engaged by high-threshold somatosensory stimuli via an ascending or descending relay that receives spinal inputs^53^.

Meanwhile, the LPBN-CM/PF pathway gets direct SDH inputs (Figure 1, S5F, G). Here, we show that the LPBN_post-spinal_ neurons terminate in the PF in addition to the CM and can play an independent role in pain processing. Moreover, we found that PF determines the effects of chronic pain on affective-motivational and somatosensory hypersensitivities (Figure 5D). In contrast, the CM is preferentially involved in the affective-motivational components of pain (Figure 5E). In the context of licking responses to chronic pain, these differences between CM and PF can be explained by the striatal projections of PF_post-LPBN_ and CM_post-LPBN_ neurons to the striatum (Figure S5). The striatal projections of the PF have been known to guide behaviors informed by sensory stimuli^22^. This finding may have implications in functional neurosurgical treatment paradigms, where targeting the PF singularly can have deterministic effects.

CIPN is known to cause mechanical allodynia^54,55^. However, we have not tested the roles of the PF_post-LPBN_ and CM_post-LPBN_ neurons in neuropathy-induced mechanical hypersensitivity. It is important to study how the PBN-CM/PF can modulate mechanical allodynia, as mechanical hypersensitivity is a common pathology in patients with neuropathic chronic pain^1,56^. Notably, mechanical hypersensitivity, as with cold, elicits shaking and guarding in mice with peripheral neuropathy and implies similar neural mechanisms underlying nocifensive behaviors in animals with cold and mechanical allodynia. Recordings from the CM-PF complex indicate the existence of mechanosensitive neurons (Figure S6C, D). In addition to CIPN, neuropathic pain in animal models induced by models such as spared-nerve injury (SNI) results in mechanical allodynia^1^ and, thus, can potentially be used to test the roles of PF_post-LPBN_ and CM_post-LPBN_ neurons.

The molecular mechanisms that facilitate hypersensitivity in the CM-PF complex post-onset of chronic neuropathic pain are not known. One of the candidates is synaptic plasticity, which has been known to be instrumental in transitioning between acute and chronic pain pathophysiology^57,58^. Recently, cell-type specific genetic manipulations perturbing synaptic plasticity in LPBN neurons have demonstrated how this molecular mechanism can play a role in consolidating learning of aversive behaviors^59^. Using similar strategies, the synaptic plasticity-related genes can be knocked out from the PF_post-LPBN_ and CM_post-LPBN_ neurons to test their role in CIPN-induced licking behaviors. The LPBN neurons are rich in neuropeptides^60^; thus, the CM-PF complex inputs can facilitate their excitability changes, as seen in mice with CIPN.

## Methods

### Experimental Model and Subject details

#### Mouse strains

Animal care and experimental procedures were performed following protocols approved by the CPSCEA at the Indian Institute of Science. The animals were housed at the IISc Central Animal Facility under standard animal housing conditions: 12 h light/dark cycle from 7:00 am to 7:00 pm with ad-libitum access to food and water; mice were housed in IVC cages in Specific pathogen-free (SPF) clean air rooms. Mice strains used: VGlut2*^Cre^* or *Slc17a6tm2*(Cre) Lowl/J(Stock number 016963); Ai14 (B6;129S6-*Gt(ROSA)26Sortm9*(CAG-tdTomato)Hze/J (Stock No 007905); Ai213 (B6;129S6-*Igs7tm213*(CAG-EGFP,CAG-mOrange2,CAG-mKate2) *Tasic*/J(Stock number 034113); BALB/cJ, Jackson Laboratories, USA. Experimental animals were between 2-4 months old.

#### Viral vectors and stereotaxic injections

Mice were anesthetized with 2% isoflurane/oxygen before and during the surgery. The craniotomy was performed using a handheld micro drill. The viral injections were administered using a Hamilton syringe (10µl) with glass-pulled needles. The viral injections (1:1 in saline) were 300nl, infused at 100nl/min. For patch clamp electrophysiological recording sections, 7- to 12-week-old male Ai213 mice were utilized for the experiment. The mice were initially anesthetized with 5% isoflurane and secured to a stereotaxic instrument (David Kopf Instruments, USA). The incision was sutured following the injection, and the mouse was placed on a warming mat until anesthesia recovery. The following were the coordinates for viral injections: LPBN (AP: −5.30 ML: +1.50 DV: −3.15); PF (AP: −2.3 ML: −0.75 DV: −3.50); CT (AP: - 1.75 ML: −0.75 DV: −3.15). The vectors used and their sources: AAV1-hSyn-Cre.WPRE.hGH (Addgene, Catalog# v126225); AAVretro-pmSyn1-EBFP-Cre (Donated by Ariel Levine, NIH); scAAV-1/2-hSyn1-FLPO-SV40p(A) (University of Zurich, Catalog# v59-1); ssAAV-9/2-hGAD1-chl-icre-SV40p(A) (University of Zurich, Catalog# v197-9); pAAV5-hsyn-DIO-EGFP (Addgene, Catalog# 50457-AAV 1); pAAV5-FLEX-tdTomato (Addgene, Catalog# 28306-PHP.S); pENN.AAV5.hSyn.TurboRFP.WPRE.RBG (Addgene, Catalog# 10552-AAV1); pAAV5-hsyn-DIO-hM4D(GI)-mCherry (Addgene, Catalog# 44362-AAV5); AAV9.syn.flex.GCaMP6s (Addgene, Catalog# pNM V3872TI-R); AAV5-hSyn-hM3D(Gq)-mCherry (Addgene, Catalog# 50474); and pAAV-Ef1a-DIO hChR2(C128S/D156A)-mCherry(Addgene, Catalog# 35504), rAAV-EF1a-DIO-Kir2.1-P2A-EGFP-WPRE-hGH (BrainVTA, Catalog# PT-1401). For rabies tracing experiments, rAAV5-EF1α-DIO-oRVG (BrainVTA, Catalog# PT-0023) and rAAV5-EF1α-DIO-EGFP-T2A-TVA (BrainVTA, Catalog# PT-0062) were injected first, followed by RV-EnvA-Delta G-dsRed (BrainVTA, Catalog# R01002) after 2 weeks. Tissue was harvested after 1 week of rabies injection for histochemical analysis. Post-hoc histological examination of each injected mouse was used to confirm that viral-mediated expression was restricted to target nuclei. The tissue was harvested one week after the rabies virus injection. Post hoc histological examination of each injected mouse was used to confirm the nuclei targeted viral expression. Stereotaxic injections to deliver viral particles in the lumbar sections were performed as previously reported^31^. 100 nl of AAV was injected at 200 µm of depth into the dorsal horn of the lumbar segments of mice under anesthesia with hamilton syringes (10µl) with glass-pulled needles.

### Fiber implantation

For fiber photometry and optogenetics, fiber optic cannulas from RWD with Ø1.25 mm Ceramic Ferrule, 200 μm Core, 0.22 NA, and L = 5 mm were implanted at the following coordinates after infusion of the viral injections: LPBN (AP:-5.30 ML:+-1.50 DV:-3.15); PF (AP: - 2.3 ML: −0.75 DV: −3.50); CT (AP: −1.75 ML: −0.75 DV: −3.15). The fibers were implanted after GCaMP6s or jRGECO1a or ChR2 infusion in LPBN, PF, or CT with light cured dental cement. Animals were allowed to recover for at least 3 weeks before performing behavioral tests. Homecage recordings were done to observe spontaneous GCaMP6s dynamics. Mice without obvious calcium dynamics were not used for the experiments. Successful labeling and fiber implantation were confirmed post hoc by staining for GFP/mCherry for viral expression and the tissue injury caused by the fiber for implantation. Animals with viral-mediated gene expression at the intended locations and fiber implantations, as observed in post hoc tests, were only included.

### Fiber Photometry

A three-channel fiber photometry system from RWD, China (R821), was used to collect data. Three light LEDs (410/470/540nm) were passed through a fiber optic cable coupled to the cannula implanted in the mouse. Fluorescence emission was acquired through the same fiber optic cable onto a CMOS camera. The photometry data was analyzed using the RWD software, and CSV files were generated. The start and end of stimuli were timestamped from the videography. All trace graphs were plotted from CSV files using GraphPad Prism 7/9 software. All heatmaps were plotted from CSV files using Python scripts. All photometry recordings under anesthesia were conducted three weeks after the GCaMP6s injection and fiber implantation.

### Fiber Photometry recording under anesthesia

A three-channel fiber photometry system from RWD, China (R821), was used to collect data. All photometry recordings under anesthesia were conducted three weeks after the GCaMP6s injection and fiber implantation. The mice were head-fixed and slightly anesthetized with a constant flow of isoflurane before the start of the recordings. During each session, the tails of the mice were pinched or touched using forceps. All sessions were videotaped from the side to capture the pinch or touch events using a camera (Logitech, USA). These event time points were used to sync with the photometry data.

### GRIN lens imaging recording and analysis

The GRIN lenses used in the experiment were 0.5mm in diameter and 5 mm in length (Inscopix, USA). Three weeks after the viral GCaMP injection, the GRIN lens was implanted on the same coordinates. A preassembled miniscope^61,62^ was purchased from LabMaker and mounted on the head under anesthesia at 1.5% isoflurane. All GRIN lens recording experiments were acquired while the mice were under anesthesia. During each session, the tails of the mice were pinched or touched using forceps. All sessions were videotaped from the side to capture the pinch or touch events using a camera (Logitech, USA). These event time points were used to sync with the cell data. The cell data acquired from these sessions was analyzed by adapting the CalmAn analysis pipeline^63^.

### Patch clamp electrophysiological preparation and recording

Virus-injected male Ai213 mice were subcutaneously injected into the left hind paw with Oxaliplatin (or saline) one day before recording. Anesthesia was induced through an intraperitoneal injection of a ketamine (100 mg/kg) and xylazine (10 mg/kg) cocktail. After confirming a negative response to the tail pinch test, the mice were transcardially perfused with ice-cold NMGD solution (composed of NMDG 93, KCl 2.5, NaH2PO4 1.25, NaHCO3 30, HEPES 20, Glucose 25, Ascorbic acid 5, Thiourea 2, Sodium pyruvate 3, MgSO4 5, CaCl2 0.5, N-acetylcysteine 12 mM, adjusted to pH 7.3 with 12N HCl). The brain was then collected and sliced into 250 μm thickness using a Compresstome VF-210-0Z (Precisionary Instruments, USA) in ice-cold NMDG solution. The brain slices were allowed to recover in 35°C warm NMDG solution for 10 minutes and subsequently incubated in room temperature aCSF (composed of NaCl 124, KCl 2.5, NaH2PO4 1.25, NaHCO3 24, HEPES 5, Glucose 12.5, MgCl2 1, CaCl2 2 mM, adjusted to pH 7.3 with 10M NaOH) for more than 30 minutes before recording.

The patch-clamp procedure was conducted using the SliceScope Pro 1000 microscope (Scientifica, USA) and PatchStar micromanipulator (Scientifica, USA). Images were captured using the Retiga Electro CCD camera (Teledyne Photometrics, USA) and Ocular software (Teledyne Photometrics, USA). The PF region was identified with DIC illumination using an 860nm LED (Thorlabs, USA) and a Plan N 4x/0.10 NA objective lens (Olympus, Japan). Neurons were detected through DIC imaging using a LUMPLFLN 40x/0.80 NA W objective lens (Olympus, Japan), and GFP fluorescence signals were detected by illuminating with a 470nm LED (Thorlabs, USA) filtered through a 49003-ET-EYFP filter cube (Chroma Technology Corporation, USA).

Thin-wall Borosilicate Glass with filament (BF150-110-10HP, Sutter Instrument, USA) was pulled using a P-1000 (Sutter Instrument, USA), adjusting the pipette resistance to 4-7 MΩ. Electrophysiological data were recorded with Axon Instruments Axon Digidata 1550B (Molecular Devices, USA), MultiClamp 700B (Molecular Devices, USA), and pClamp 11 software (Molecular Devices, USA) at a 2 kHz sampling rate.

Voltage clamp recording was conducted at −70 mV using a Cs-gluconate-based internal solution (composed of D-gluconic acid 110, CsOH 110, CaCl2 0.1, MgATP 4, Na2GTP 0.4, Na^2^ Phosphocreatine 10, EGTA 1.1, HEPES 10, TEA-Cl 8 mM). Miniature EPSCs were recorded under channel blockers, including Tetrodotoxin (Hello Bio, USA) 1 μM, Picrotoxin 100 μM, and Bicuculline 10 μM. Recordings were made over 3 minutes, and the events were detected and analyzed using Clampfit software (Molecular Devices, USA). Before analysis, the data were low-pass filtered below 0.2 kHz.

Light-evoked EPSCs were recorded under the aCSF with Picrotoxin 100 μM and Bicuculline 10 μM. 470 nm LED stimulation was delivered through the 40x objective lens, with 1.2 mW of light intensity at the sample plane, for a 5 ms duration. The light stimulation was performed at least 10 times at 10-second intervals, and the amplitude of the first 10 trials was measured and averaged. The neurons with extremely large jitter were excluded from the analysis to exclude polysynaptic response.

### Behavioral assays

A single experimenter handled behavioral assays for the same cohorts. The experimenter was blinded to the identity of the animals. The mice were habituated in their home cages for at least 30 minutes in the behavior room before starting the experiment. An equal male-to-female ratio was maintained in every experimental cohort and condition unless otherwise stated, with no significant differences seen between sexes in the responses recorded from the behavioral experiments. Wherever possible, efforts were made to keep the usage of animals to a minimum. Oxaliplatin (2mg/ml) was injected into the left paw, and the mice were left in their home cages in the behavior room for a minimum of 4 hours before any behavioral experiments to cause persistent neuropathic pain and thermal hypersensitivity^15^. For all the behavioral experiments, the first set of experiments was performed 4 hours after oxaliplatin injection. The second set of experiments was done within 12-14 hours to retain the maximum cold hypersensitivity caused by oxaliplatin-induced peripheral neuropathy.

Additionally, for every cohort, half the mice had baseline experiments in the first set and manipulation experiments in the second set, while for the other half, the sequence of experiments was reversed to control for any biases that may exist in responses across time. For chemogenetic experiments, Deschloroclozapine (100 µg/kg) dissolved in water and injected intraperitoneally at least 30 minutes before behavioral experiments. For optogenetic experiments, the fiber-coupled lasers (RWD, China) were connected to the optic ferrules implanted on the heads of mice. The mice were left for 20 minutes to habituate to the fiber and placed in their respective chamber depending on the behavioral assay. During experiments, the mice received photostimulation with a power of 20mW at 15Hz with a pulse width of 5ms.

### Thermal plate experiments

According to the manufacturer’s instructions, we used a hot-cold analgesia plate from Orchid Scientific for these experiments^64,65^. We placed mice on the thermal plate covered by an acrylic box at various temperatures to perform three assays. The gradient cold plate assay lasted 5 minutes, where the temperature dropped from 24 to 0 degrees at 4.8 degrees/minute. The static cold plate assay lasted for 5 minutes, where the temperature was set to 12 degrees. The static hot plate assay lasted for 45 seconds, where the temperature was set to 52 degrees. All the experiments were videotaped simultaneously with three wired cameras (Logitech, USA) placed horizontally and scored manually post hoc. All experiments were scored by an experimenter blinded to the different experimental conditions.

### Real-time place aversion test

This assay used a custom-built box with two identical 20 x 20cm chambers with a middle partition to observe avoidance responses during neural stimulation. On day 1, the mice were placed in one chamber with the fiber for optical stimulation attached to the fiber implanted in their heads. They were allowed to explore both chambers without optogenetic stimulation. On day 2, one chamber was the control with no optogenetic stimulation, and the other was the stimulation chamber. Each day, the assay was run for a total of 20 minutes. The experiments were videotaped using a camera (Logitech, USA) from the top to track the mice across chambers using DLC.

### Conditioned place aversion test

For this assay, a custom-built box with two 32 x 32cm chambers connected by an 11 x 8.5 cm long middle passage was used. One chamber had black and white striped walls, while the other had black walls. The mice could explore both chambers without restrictions on the pre-training day (day 0). On the first training day (day 1), the mice were injected with DCZ 20 minutes before being placed in the middle passage, and access to the black chamber was restricted, allowing access only to the striped chamber. On the second training day (day 2), the mice were injected with saline 20 minutes before being placed in the middle passage, and access to the striped chamber was restricted, allowing access only to the black chamber. This was repeated for four more days, and on the post-training day (day 7), the mice were placed in the middle passage with free access to both chambers to observe if they displayed avoidance towards any one chamber. Each day, the test was run for a total of 30 minutes. The experiments were videotaped using a camera (Logitech, USA) from the top to track the mice across chambers using DLC.

### Temperature place avoidance test

The dual thermal analgesia plate was purchased from Orchid Scientific and was used according to the manufacturer’s instructions. The plate has two 20 x 20cm sides, which can be set to different temperatures. One side of the dual thermal plate was set to 24 degrees and the other to 4 degrees. Both the plates were covered by an acrylic box with a partition in the middle to allow mice free access to both sides. Mice were placed on the 24-degree side. The test was run for a total of 20 minutes. The experiments were videotaped using a camera (Logitech, USA) from the top to track mice across the chambers, and the videos were scored post hoc by an experimenter blinded to the different experimental conditions for time spent on each side.

### Immunostaining

Mice were anesthetized with isoflurane and perfused intracardially with 1X Phosphate Buffered Saline (PBS) (Takara, Japan) and 4% Paraformaldehyde (PFA) (Ted Pella, Inc., USA), consecutively for immunostaining experiments. The brains were harvested and further fixed in 4% PFA overnight at 4°, then stored in 30% sucrose overnight. The brains were embedded in an OCT medium and were cut at a thickness of 50µm sections using a cryostat (RWD). Brain sections were floated in PBS and mounted onto glass slides before staining.

Tissue sections were rinsed in 1X PBS and incubated in a blocking buffer (2% Bovine Serum Albumin (BSA); 0.3% Triton X-100; PBS) for 1 hour at room temperature. Sections were incubated in primary antibodies overnight in a blocking buffer at room temperature. Sections were rinsed 1-2 times with PBS and incubated for 2 hours in Alexa Fluor conjugated goat anti-rabbit/ chicken or donkey anti-goat/rabbit secondary antibodies (Invitrogen) at room temperature, washed in PBS, and mounted in VectaMount permanent mounting media (Vector Laboratories Inc.) onto charged glass slides (Globe Scientific Inc.). We used an upright fluorescence microscope (Khush, Bengaluru) (2.5X, 4X, and 10X lenses) and ImageJ/FIJI image processing software to image and process images to verify the anatomical location of cannulae implants. For the anatomical studies, the images were collected with 10X and 20X objectives on a laser scanning confocal system (Leica SP8 Falcon, Germany) and processed using the Leica image analysis suite.

### Biocytin visualization

Internal solutions containing 0.5% biocytin were used to visualize the recorded neurons during recordings. After recording, slices were incubated in 4% paraformaldehyde (PFA) in 0.1 mM PBS at 4 degrees Celsius for 1-2 nights. The slices were washed three times with 0.1 mM PBS and incubated in 0.3% Triton-X in 0.1 mM PBS with 1/500 concentrated Alexa-647 conjugated Streptavidin (Thermo Fisher Scientific, USA) at 4 degrees Celsius overnight. After three washes with 0.1 mM PBS, the slices were mounted with DAPI Fluoromount-G (Southern Biotech, USA).

### Deep lab cut tracking

DeeplabCut (DLC) was used to track the movement of mice under different conditions compared to the recorded videos during the RTPA and CPA assays. A total of 1200 frames were labeled across 12 videos were used for the training dataset. Multiple body points were labeled, including the tail, spine, and nos,e but only the nose point was used to track mice movement. The labeling and using the nose point were performed as previously reported^64^.

### Statistical analysis

The statistical analysis utilized included t-tests to compare two sets of data and one-way ANOVA for multiple data sets. All statistical analyses were performed using GraphPad Prism 7/9 software. ns>0.05, *<0.05, **<0.01, ***<0.001, ****<0.0001.

## Abbreviations

LPBN: Lateral Parabrachial Nucelus
CM: Centro Median nucleus
PF: Parafascicular nucelus
RVM: Rostral Ventromedial Medulla
CeA: Central Amygdala
LH: Lateral hypothalamus
Str: Striatum
Fr: Fasciculus retroflexus
CIPN: Chemotherapeutic drug-induced peripheral neuropathy
ILN: Intralaminar thalamus
SDH: Spinal Dorsal Horn
Lum: Lumbar spinal cord
ChR2: Channelrhodopsin-2
mEPSC: Miniature Excitatory Postsynaptic Currents
EPSC: Excitatory Postsynaptic Currents
RTPA: Real-time place aversion
CPA: Conditioned place aversion test
TPP: Temperature plate preference
DCZ: Deschloroclozapine
AUC: Area under the curve

## Acknowledgement

We thank the DST-SERB (CRG/2021/1005124), IndiaAlliance Intermediate Fellowship (IA/I/19/2/504640), and IISc Bengaluru for funding our research.

---

**Figure S1.**
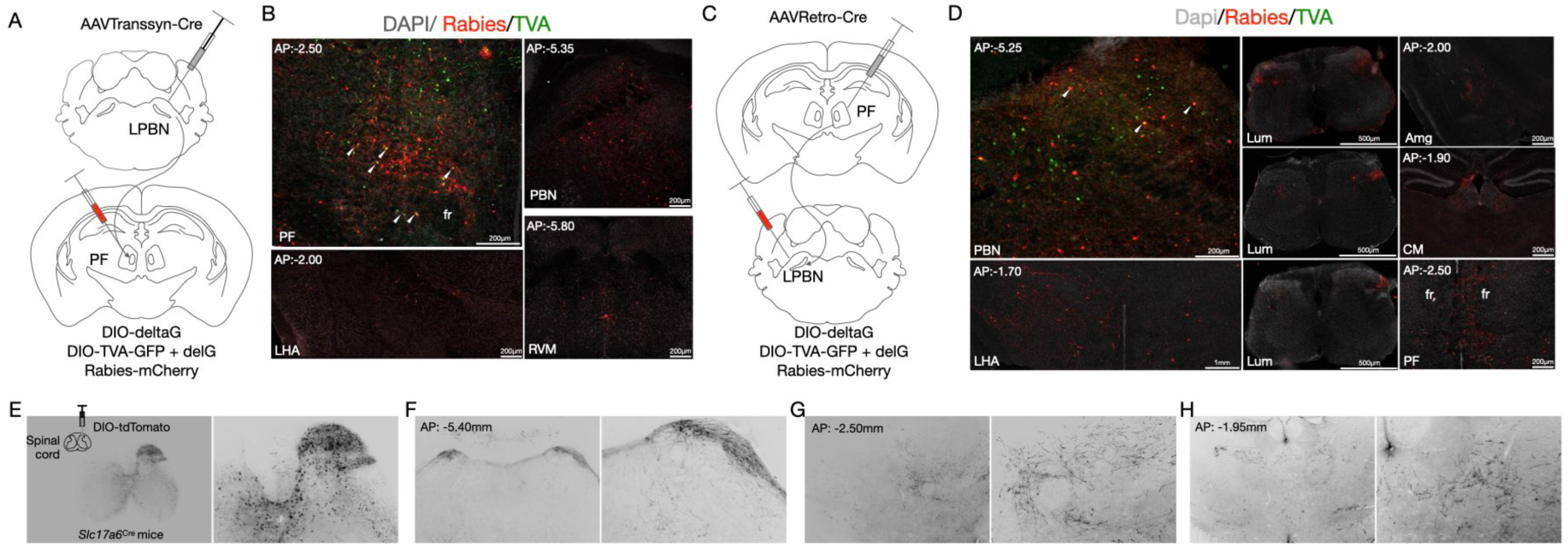
(a) The intersectional genetic strategy used for tracing the pre-synaptic partners of the PF_post-LPBN_ neurons with retrograde monosynaptic rabies viral tracing. AAVTrannsyn-Cre was injected in the LPBN, AAV5-EF1α-DIO-oRVG, and rAAV5-EF1α-DIO-EGFP-T2A-TVA were injected in the PF, followed by RV-EnvA-ΔG-dsRed after 3 weeks. (b) Representative confocal image of a coronal section of the PF (Top left image) showing expression of rabies (red) and TVA (green) expression in PF_post-LPBN_ cells along with the starter cells (yellow cells). A representative confocal image of coronal sections of LPBN (top right image), lateral hypothalamic area (LHA) (bottom left image), and RVM (bottom right image) shows the expression of retrograde rabies cells (red). (c) The strategy for tracing the inputs that LPBN cells receive upstream of the PF using retrograde rabies tracing. Receives input from the PBN cells using retrograde rabies tracing. AAVretro-pmSyn1-EBFP-Cre was injected in the PF, rAAV5-EF1α-DIO-oRVG, and rAAV5-EF1α-DIO-EGFP-T2A-TVA were injected first, followed by RV-EnvA-ΔG-dsRed after 3 weeks in the LPBN. (d) Representative confocal image of a coronal section of the LPBN (Top left image) showing expression of rabies (red) and TVA-GFP (green) expression in LPBN upstream of the PF along with the starter cells (yellow cells). Representative confocal image of coronal sections of LHA (Bottom left image), spinal cord sections (Centre panel), CeA (Right top image), CM (Right middle image), and PF (Right bottom image) showing the expression of retrograde rabies cells (red). (e) The strategy used to label the projections of the spinal dorsal horn neurons (Top right). AAV5-hSyn-DIO-tdTomato was injected in the spinal cord of *Slc17a6^Cre^* mice. A representative confocal image of a coronal section of the spinal cord 10x (Right image) and magnified 20x (Left image). (f) A representative confocal image of a coronal section of the LPBN; (g) CM; and (h) PF, 10x (Right images) and magnified 20x (Left images).

**Figure S2.**
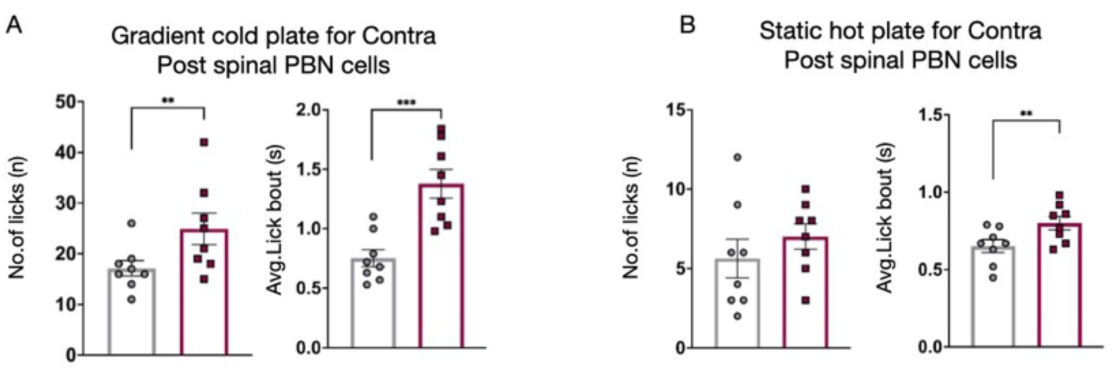
(a) Quantification of lick frequency (17.13±1.54 compared to 24.88±3.11, respectively; t-test, \*\**P*=0.005) and length of the lick bouts (0.75±0.07 compared to 1.38±0.12, respectively; t-test, \*\*\**P*=0.0004) on the gradient cold plate assay (24 to 0°C over 5mins) with an i.p injection of saline or an i.p injection of DCZ to excite the hM3Dq-expressing LPBN_post-spinal cord_ cells (n=8). (b) Quantification of lick frequency (5.63±1.21 compared to 7.00±0.80, respectively) and length of the lick bouts (0.65±0.04 compared to 0.80±0.04, respectively; t-test, \*\**P*=0.006) on the static hot plate assay (52°C over 45s) with an i.p injection of saline or an i.p injection of DCZ to excite the LPBN_post-spinal cord_ neurons (n=8).

**Figure S3.**
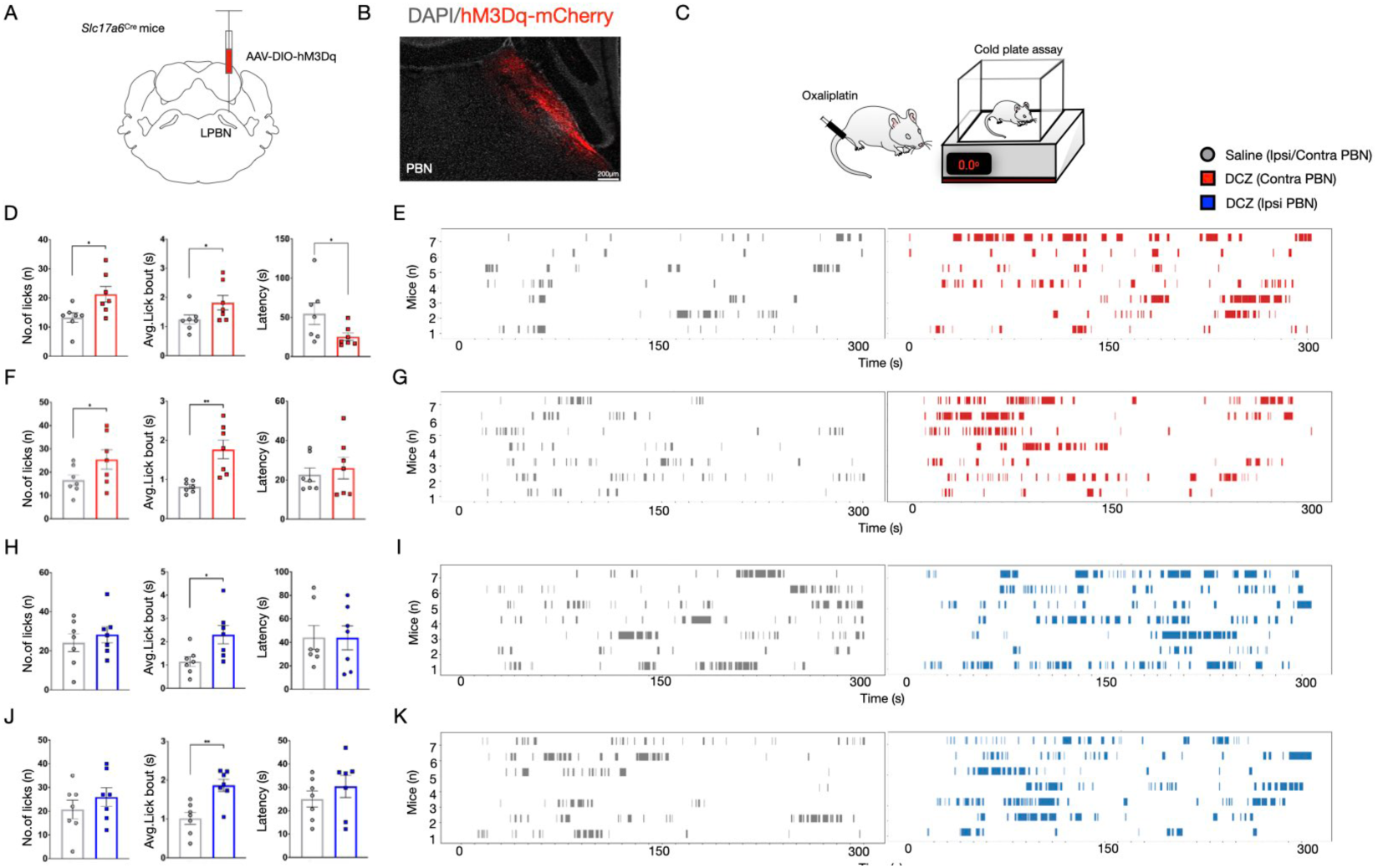
(a) Diagram representing the strategy used to excite the LPBN neurons. AAV5-hSyn-hM3D(Gq)-mCherry was stereotaxically injected into the LPBN in *Slc17a6^Cre^* mice. (b) Representative confocal images of the coronal section of the LPBN showing expression of the mCherry tagged hM3Dq (red). (c) Schematic of the intraplantar injection of oxaliplatin or saline (control) and the mice placed in a thermal plate for the cold or hot plate assay. (d) Quantification of lick frequency (13.23±1.60 compared to 21.29±2.65, respectively; t-test, \**P*=0.02), length of the lick bouts (1.24±0.15 compared to 1.82±0.25, respectively; t-test, \**P*=0.01) and latency to lick (54.74±13.69 compared to 25.33±4.74, respectively; t-test, \**P*=0.02) on the gradient cold plate assay (24 to 0°C over 5mins) with an i.p injection of saline or an i.p injection of DCZ to excite the contra-lateral LPBN (to the oxaliplatin-injected paws) (n=7). (e) Raster plots showing licking across a single session of gradient cold plate assay with saline (left plots) or DCZ to excite the contra LPBN (right plots) (n=7). (f) Quantification of lick frequency (16.57±2.24 compared to 25.43±4.15, respectively; t-test, \**P*=0.04), length of the lick bouts (0.89±0.06 compared to 1.77±0.24, respectively; t-test, \*\**P*=0.005) and latency to lick (22.64±3.40 compared to 26.01±5.53, respectively) on the static cold plate assay (12°C over 5mins) with an i.p injection of saline or an i.p injection of DCZ to excite the contra-LPBN (n=7). (g) Raster plots showing licking across a single session of static cold plate assay with i.p. saline (left plots) or DCZ the contralateral LPBN neurons (right plots) (n=7).

**Figure S4.**
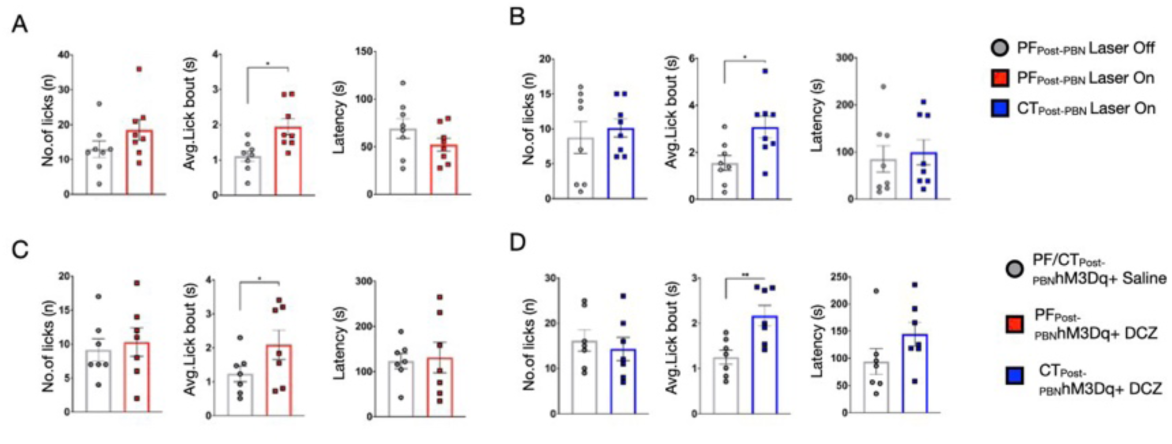
(a) Strategy used to label the PF_post-LPBN_ and CM_post-LPBN_ neurons with GFP and tdTomato, respectively. AAVTrannsyn-Cre was stereotaxically injected in the LPBN and AAV5-hsyn-DIO-EGFP in the PF and AAV5-hSyn-DIO-tdTomato in the CM. (b) Representative confocal images of the coronal sections of the CM (Left image) showing expression of TdTomato (red) and the PF (right image) showing expression of GFP (green). (c) Representative confocal images of the coronal section of the LPBN (left image), RVM (Centre image), and Striatum (right image) showing expression of TdTomato (red) and GFP (green). (d) Representative confocal images of the coronal sections of the striatum arraigned from the posterior to the anterior stereotaxic coordinates (left to right).

**Figure S5.**
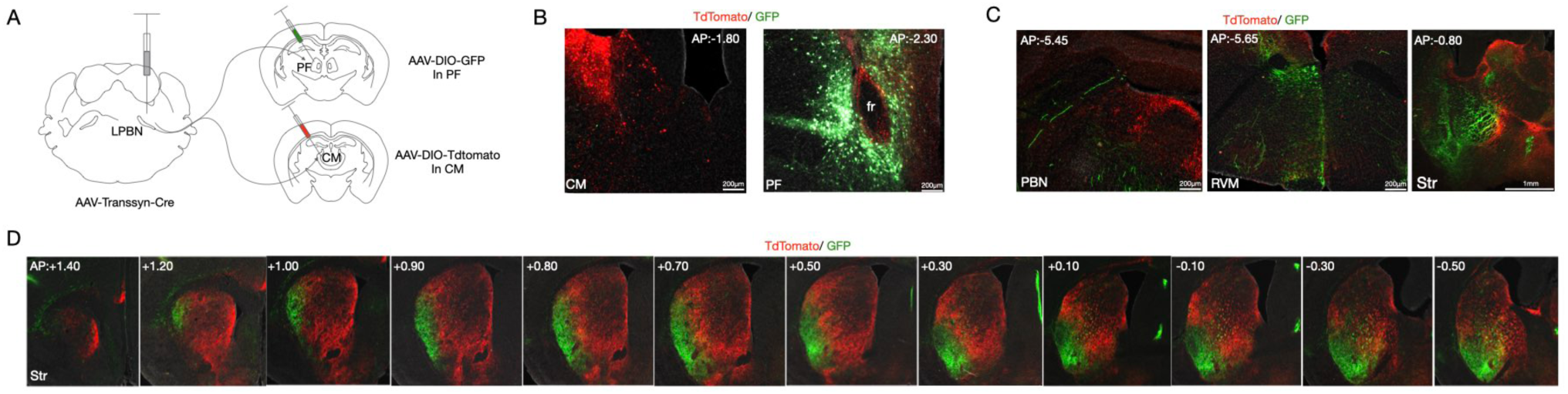
(a) The strategy used to express GCaMP6s in the PF_post-LPBN_ neurons in wild-type mice with the transsynaptic viral strategy. (b) A representative confocal image of the coronal section of the PF shows the expression of GCaMP6s in the PF_post-LPBN_ neurons. (c) Fluorescence change in the PF_post-LPBN_ neurons as measured in dF/F for mechanical low threshold (brush) and high threshold (pinch) stimuli in mice with CIPN, demonstrating mechanical allodynia and hyperalgesia under anesthesia. (d) Quantification comparing the average GCaMP6s activity (AUC) during low threshold (brush) and high threshold (pinch) stimuli in mice with CIPN (3.08±0.26 compared to 7.64±0.70, respectively; t-test, \*\*\*\**P*<0.0001) in the PF_post-LPBN_ neurons (n=3). (e) A diagrammatic representation of the strategy used to express GCaMP6s in the PF neurons in *Slc17a6^Cre^* mice (e) Fiber photometry recordings show that the contralateral PF_post-LPBN_ neurons are preferentially engaged while the mice lick due to cold allodynia. (f) A representative confocal image of the coronal section of the PF shows the expression of GCaMP6s in the *Slc17a6*-expressing PF neurons. (f) Quantification comparing the average GCaMP6s activity (AUC) during ipsilateral and contralateral licks (16.01±3.98 compared to 37.52±6.68, respectively; t-test, \*\**P*=0.001) in the *Slc17a6*-expressing PF neurons (n=2).

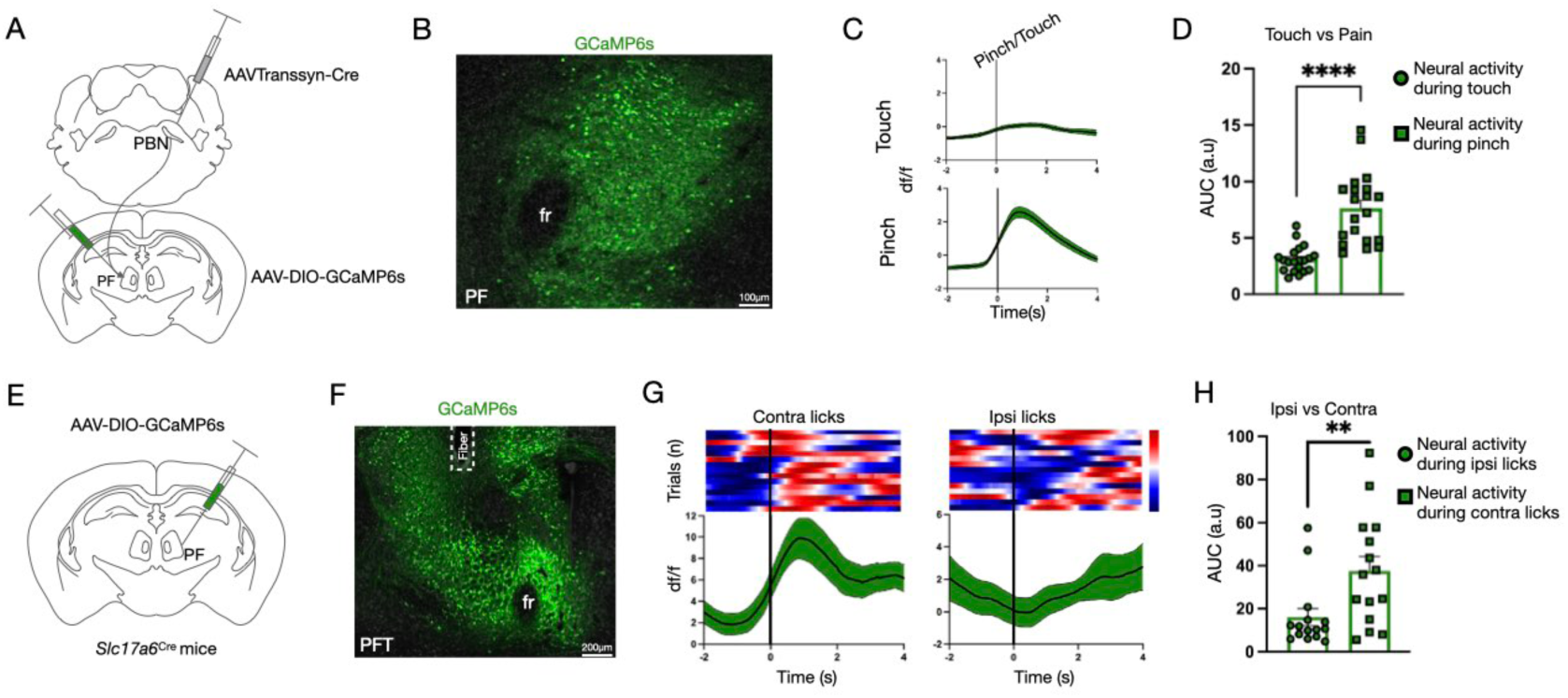

